# Early plasma proteomic alterations precede amyloidosis diagnosis, reflecting cardiac and immune dysregulation

**DOI:** 10.64898/2026.05.08.723738

**Authors:** Seyed Alireza Hasheminasab, Mohammad H. Kazeroun, Joshua Fieggen, Lei Clifton, Berrin Balik, Devi Nandana Suchitra Devi, Jesslyn Choo, Austeja Bakulaite, Udo Oppermann, Nikant Sabharwal, Karthik Ramasamy, Ashutosh D. Wechalekar, Anjan Thakurta

## Abstract

Systemic amyloidosis is typically diagnosed only after irreversible organ damage has occurred, limiting the effectiveness of available therapies. Whether the disease is preceded by detectable molecular changes long before clinical presentation has remained unclear. Here, we leveraged population-scale plasma proteomics and longitudinal follow-up from the UK Biobank to investigate early circulating protein signatures associated with future diagnosis of amyloidosis. Among approximately 53,000 participants with proteomic profiling, we identified 61 individuals who developed amyloidosis up to 14 years after protein assessment. Differential expression and correlation analyses identified a seven-protein panel, including MYL3, MYBPC1, NT-proBNP, NPPB, FCRLB, IGFBP1, and FABP1, consistent with early cardiac stress and immune dysregulation. Time-to-event modelling demonstrated robust stratification of amyloidosis risk and timing. Importantly, a parsimonious subset of these proteins retained strong predictive performance, indicating that a reduced set of biologically informative markers is sufficient for risk stratification. Furthermore, these proteomic signals were not explained by pre-existing cardiac disease, clonal haematopoiesis, or related plasma cell disorders, indicating that they capture disease-specific biological processes preceding clinical diagnosis. Together, these findings show that amyloidosis is preceded by persistent plasma proteomic alterations, providing a framework for early risk stratification and insight into the preclinical biology of this under-recognised disease.

## Introduction

Systemic amyloidosis comprises a heterogeneous group of protein misfolding disorders characterised by the extracellular deposition of insoluble amyloid fibrils in tissues and organs, leading to progressive organ dysfunction and substantial morbidity and mortality. The clinical manifestations of amyloidosis vary widely depending on the precursor condition and the organs involved, but prognosis is mainly dominated by cardiac and renal involvement. Among systemic forms, immunoglobulin light-chain (AL) amyloidosis and transthyretin (ATTR) amyloidosis account for the majority of cases. AL amyloidosis typically follows an aggressive clinical course driven by toxic light-chain production from clonal plasma cells, whereas ATTR amyloidosis, particularly in its wild-type form, often presents later in life with predominant cardiac involvement. Despite major therapeutic advances, overall outcomes remain poor for both disease subtypes, mainly because diagnosis often occurs only after irreversible organ damage has developed.

Delayed diagnosis is an important characteristic of amyloidosis and a major contributor to poor outcomes. In its early stages, the disease is typically clinically silent or presents with non-specific signs and symptoms causing delayed diagnosis, missed opportunities for intervention, and prolonged diagnostic pathways. Advanced cardiac involvement at diagnosis is associated with severely reduced survival time in AL amyloidosis and its delayed recognition often causes progressive heart failure that is resistant to treatment. Current diagnostic and prognostic tools largely reflect established organ dysfunction rather than early disease-associated biological signals. Cardiac biomarkers such as N-terminal pro–B-type natriuretic peptide (NT-proBNP) and troponins are essential in risk stratification and disease staging^1,2,3^ but considered only after organ involvement has already developed. As a result, instead of utilising biomarkers to identify high-risk individuals before emergence of symptoms or irreversible organ injury, they are primarily used once clinical suspicion and overt symptoms have already arisen.

Emerging epidemiological data also suggest that amyloidosis is substantially under-recognised in the general population. Wild-type ATTR amyloidosis, considered a rare disorder, is expected to be identified among individuals with heart failure with preserved ejection fraction (HFpEF), a common and heterogeneous clinical syndrome. Autopsy and biopsy-based studies reported myocardial amyloid deposits in approximately 14 percent of older patients with HFpEF^4^, highlighting the potential under-recognition of cardiac amyloidosis in clinical practice. Recent studies indicated that targeted screening for organ damage biomarkers in monoclonal gammopathy of undetermined significance (MGUS) can identify a fraction of individuals with presymptomatic AL amyloidosis^5^. However, these strategies rely on relatively broad screening filters that detect only a limited proportion of patients with presymptomatic disease within the enriched MGUS population. These observations highlight the persistent challenge of delayed amyloidosis diagnosis, underscoring the need for techniques capable of detecting disease at an earlier, clinically silent stage in the general population.

Despite this need, early detection of amyloidosis has historically been challenging. The rarity of clinically diagnosed disease, combined with long and variable latency periods, has limited systematic investigation of presymptomatic stages. Most prior investigations have been cross-sectional or focused on patients at diagnosis time, limiting the understanding of potential events that precede overt disease. Although recent studies have applied artificial intelligence to electrocardiography, echocardiography, imaging, and routine laboratory data to enhance detection of cardiac amyloidosis, these approaches are largely cross-sectional and are typically applied after clinical suspicion has already emerged^6,7,8,9^.

Advances in high-throughput plasma proteomic profiling have enabled simultaneous measurement of thousands of circulating proteins from small volumes of plasma, leading to a comprehensive view of systemic biology. These technologies enable systematic investigation of molecular alterations preceding clinical disease in large population cohorts with long-term follow-up. Population-scale plasma proteomics in the UK Biobank has recently been shown to modestly improve prediction of common cardiovascular outcomes and to identify dysregulated immune proteins up to a decade before clinical diagnosis in multiple myeloma^10,11^. However, whether such approaches can predict rare diseases with long preclinical phases, such as amyloidosis, remains largely unexplored.

With plasma proteomic information available for approximately 50,000 participants and up to two decades of linked follow-up health records, the UK Biobank enables investigation of incident amyloidosis in individuals who were clinically unaffected at the time of protein measurement. This design allows interrogation of circulating proteins years before diagnosis, disentangling early disease-associated biology from downstream consequences of organ dysfunction or treatment.

In this study, we leveraged population-scale plasma proteomics from the UK Biobank to investigate circulating protein signatures associated with future amyloidosis diagnosis years before clinical manifestation. We identified a seven-protein panel comprising MYL3, MYBPC1, NT-proBNP, NPPB, FCRLB, IGFBP1, and FABP1 that is associated with future amyloidosis diagnosis years before clinical presentation. Using time-to-event modelling and explainable machine-learning approaches, we demonstrate that a parsimonious subset of these proteins enables robust prediction of both risk and timing of disease over extended preclinical intervals. These findings reveal early cardiac and immune alterations that precede clinically apparent disease and provide a framework for early amyloidosis risk stratification.

## Results

### Cohort derivation and study design

To investigate whether circulating plasma proteins can predict the future development of amyloidosis, we analysed participants from the UK Biobank with available plasma proteomic profiling and longitudinal health record follow-up (Fig. 1a). Plasma protein measurements were recorded for almost 10% of the UKB population at a baseline assessment using a high-throughput proximity extension assay platform, and this timepoint was marked as the index date for all subsequent analyses.

**Figure 1.**
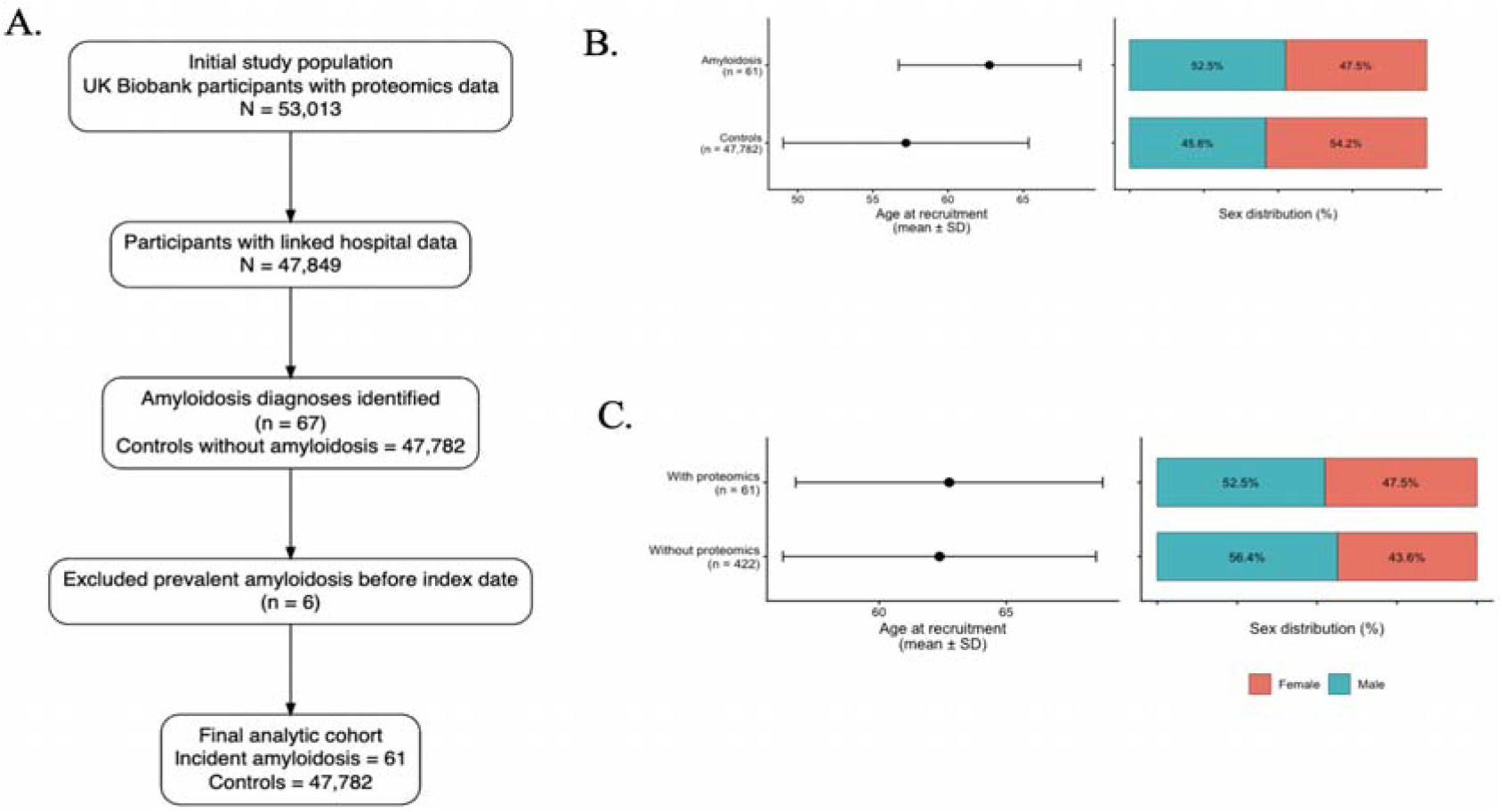
Cohort derivation and study population characteristics. (a) Flow diagram showing derivation of the study population from the UK Biobank proteomics cohort. (b) Baseline characteristics of participants stratified by incident amyloidosis status, showing age and sex distribution. (c) Comparison of amyloidosis cases with and without available proteomic data, demonstrating that the proteomics subset is representative of the broader amyloidosis population.

Among participants with proteomic data, amyloidosis cases were identified using linked electronic health records from hospital inpatient data. Cases were identified using International Classification of Diseases (ICD-10) codes corresponding to systemic amyloidosis. Individuals with a recorded diagnosis of amyloidosis prior to the index date were excluded to ensure that only incident cases were analysed. The temporal distribution of diagnosis and related clinical events is shown in Supplementary Fig. S4.

The final study population covered 61 individuals who developed amyloidosis after plasma protein assessment and 47,782 controls who remained amyloidosis free during follow-up according to their diagnosis records. The median time from plasma protein measurement to amyloidosis diagnosis was approximately 9 years, with a range extending up to 14 years. This enabled investigation of proteomic alterations during the extended preclinical phase of disease.

Baseline demographic characteristics of amyloidosis cases and controls were broadly comparable (Fig. 1b), although cases were older at the time of protein assessment (mean age of approximately 63 years), consistent with the known age distribution of amyloidosis. Sex distribution was similar between groups. The demographic characteristics of amyloidosis cases with available proteomic data were comparable to those of amyloidosis cases in the wider UK Biobank without proteomic profiling (Fig. 1c), supporting the representativeness of the analysed subset. Baseline characteristics of the full cohort and the proteomics subset are shown in Supplementary Tables S1 and S2, and the overlap of amyloidosis-related comorbidity patterns between participants with and without proteomic data is shown in Supplementary Fig. S1. All predictive modelling analyses were performed using the subset of participants with available proteomic data.

This study design, combining a single baseline proteomic measurement with extended longitudinal follow-up and incident disease ascertainment, enables systematic investigation of early circulating protein signatures associated with the future development of amyloidosis in a large population cohort.

### Early plasma proteomic alterations precede clinical amyloidosis diagnosis

To characterise early circulating protein changes associated with the future amyloidosis diagnosis, we first performed an unsupervised exploration of the plasma proteomic profiles. Principal component analysis (PCA) of the full proteomic dataset showed substantial overlap between amyloidosis cases and controls, with no clear global separation between groups (Fig. 2a). This indicates that the disease-associated signal is not driven by large-scale global variation in overall protein expression but rather reflects the coordinated alterations across a subset of proteins.

**Figure 2.**
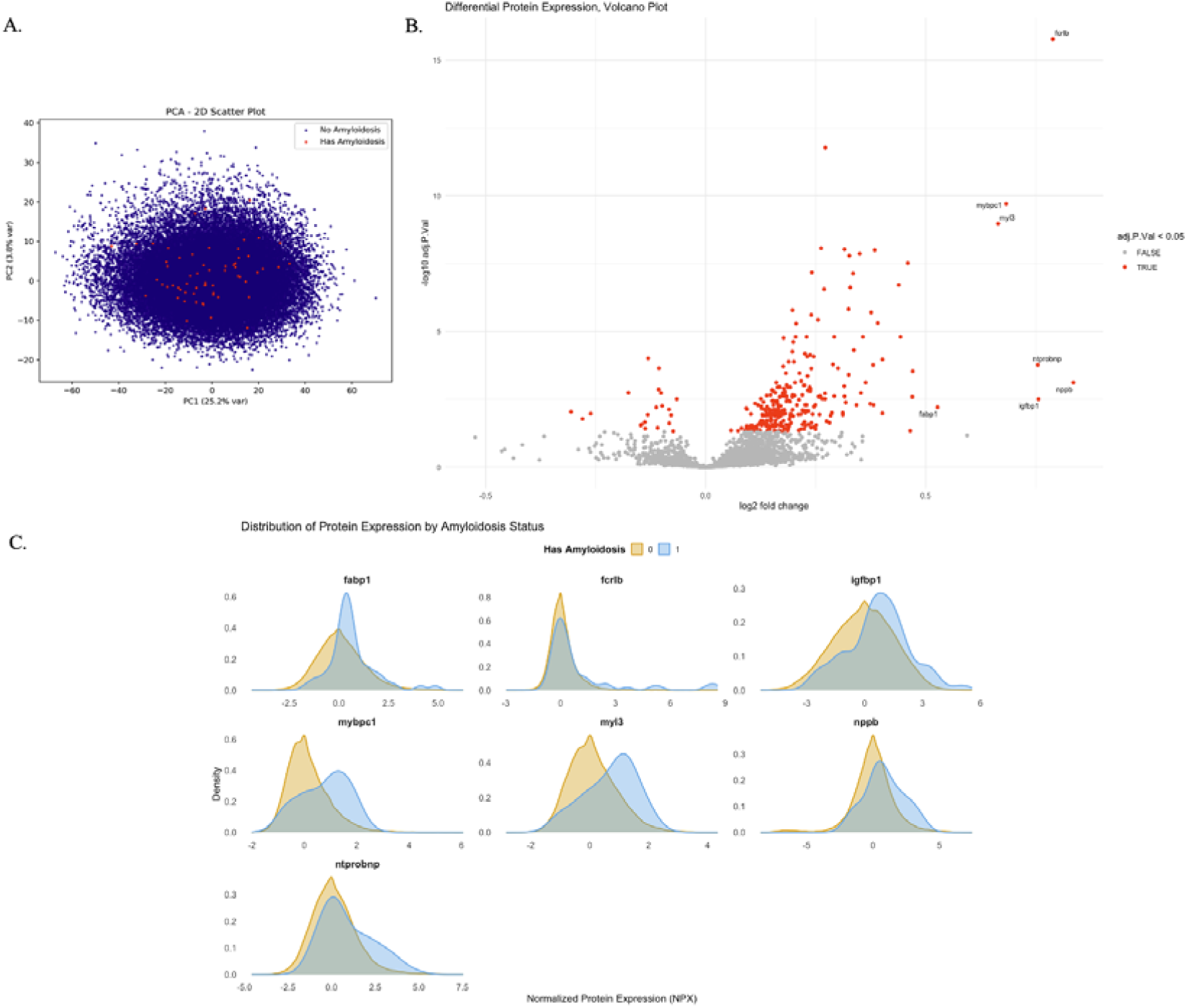
Plasma proteomic alterations preceding amyloidosis diagnosis. (a) Principal component analysis of plasma proteomic profiles in individuals who developed amyloidosis and controls, demonstrating substantial overlap and indicating that disease-associated differences are not driven by global proteomic variation. (b) Differential protein expression analysis comparing amyloidosis cases and controls using imputed proteomic data. Seven significantly differentially expressed proteins are highlighted. Age- and sex-adjusted analyses are shown in Supplementary Fig. S2. (c) Distribution of selected plasma proteins in amyloidosis cases and controls. Despite overlap between groups, a systematic shift in expression is observed in cases during the preclinical phase.

We next performed differential protein expression analysis comparing amyloidosis cases with controls using linear models with empirical Bayes moderation. Multiple proteins were significantly differentially expressed, including several cardiac-associated markers such as MYL3, MYBPC1, NPPB, and NT-proBNP, as well as immune- and metabolic-related proteins including FCRLB, IGFBP1, and FABP1 (Fig. 2b). Differential expression analysis was performed using imputed proteomic data to account for missing values (Fig. 2b), while age- and sex-adjusted analyses yielded similar but attenuated effects (Supplementary Fig. S2). These proteins showed consistent directional changes and remained significant after adjustment for age and sex, with the exception of FABP1, which was attenuated. Given the limited sample size and its potential biological relevance, FABP1 was retained for downstream analyses, including predictive modelling, where age and sex were explicitly accounted for.

To further characterise these differences, we examined the distribution of key proteins in cases and controls. Although there was substantial overlap at the population level, individuals who later developed amyloidosis showed a consistent shift toward higher expression levels for several proteins (Fig. 2c). In some individuals, elevations were detectable many years before diagnosis, indicating that early pathological processes are already reflected in the circulating proteome.

These findings demonstrate that amyloidosis is preceded by reproducible but subtle alterations in a defined subset of proteins. While not separable at the global proteome level, these changes are detectable through targeted differential expression analysis and provide a biologically plausible candidate set for downstream modelling. This set served as the biological foundation for subsequent modelling and interpretability analyses.

### Preclinical amyloidosis is characterised by coordinated cardiac and immune proteomic signatures

To further interpret the biological basis of the observed proteomic alterations, we examined relationships among the key differentially expressed proteins and their expression patterns across individuals.

Correlation analysis revealed strong co-expression among cardiac-associated proteins, including MYL3, MYBPC1, NPPB, and NT-proBNP, consistent with a shared biological origin related to myocardial stress and structural remodelling (Fig. 3a). These proteins formed a strongly correlated group, suggesting activation of cardiac-related pathways long before clinical diagnosis.

**Figure 3.**
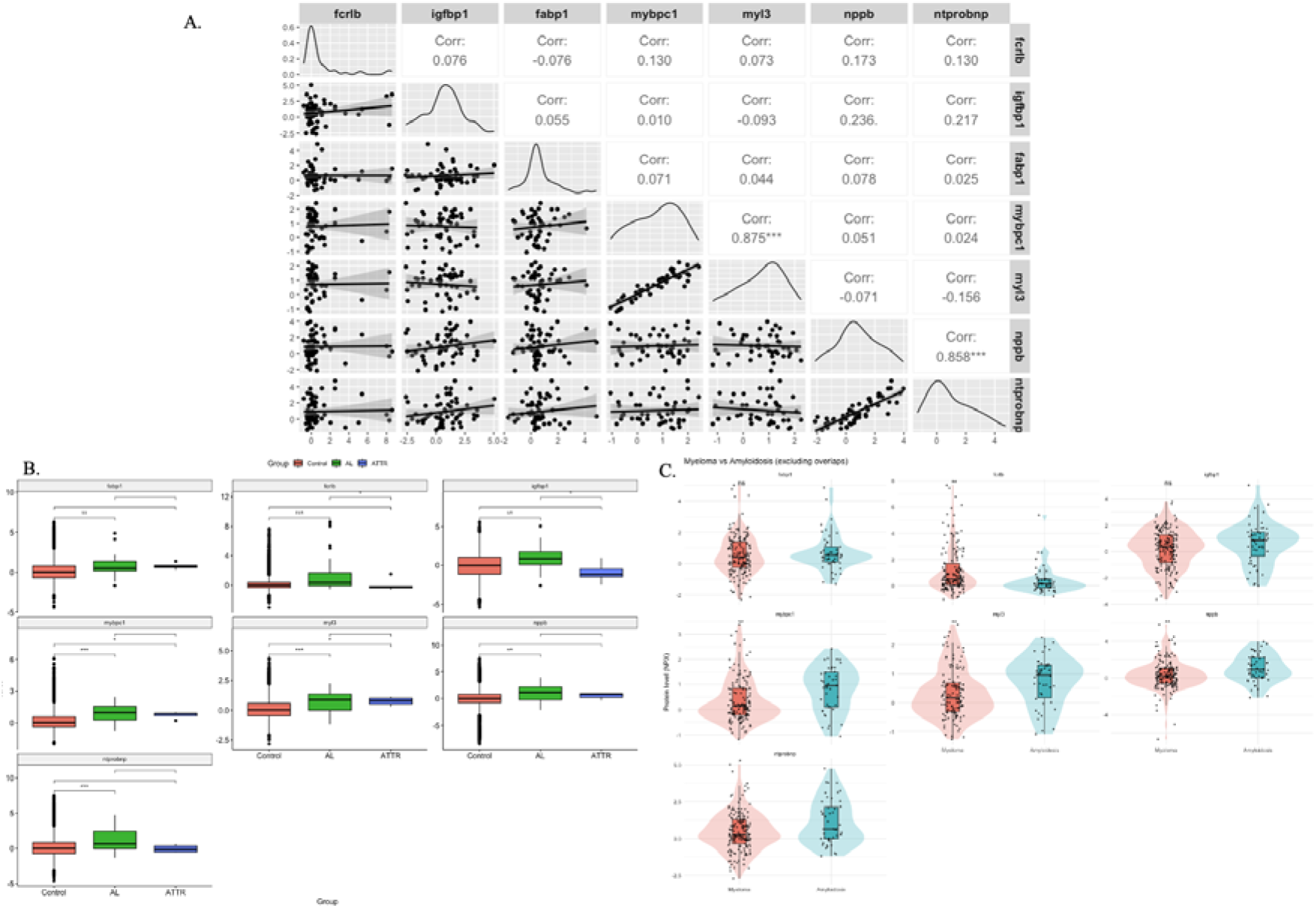
Early amyloidosis is associated with linked cardiac and immune proteomic signatures. (a) Correlation matrix of selected plasma proteins demonstrating clustering of cardiac-associated markers and weaker correlations with immune and metabolic proteins. (b) Distribution of selected proteins across amyloidosis subtypes, showing preservation of the proteomic signal in AL amyloidosis. (c) Comparison of key plasma proteins between amyloidosis and multiple myeloma cohorts, highlighting both shared and disease-specific proteomic features.

In contrast, immune-related proteins, particularly FCRLB, showed weaker correlations with cardiac proteins, indicating a partially independent biological axis. Additional proteins such as IGFBP1 and FABP1 showed weak-to-moderate correlations, suggesting involvement in broader systemic metabolic or inflammatory processes. Together, these findings indicate that amyloidosis is preceded by alterations involving cardiac, immune, and metabolic pathways rather than isolated organ-specific pathology.

To place these protein-level relationships into a broader biological context, we next performed pathway enrichment analysis. Pathway enrichment analysis of age- and sex-adjusted proteomic associations revealed enrichment of extracellular matrix organisation, cardiac and muscle development pathways, and immune-related processes including complement activation, coagulation, and inflammatory response (Supplementary Fig. S6). These findings support early cardiac structural remodelling alongside systemic immune activation.

These pathway and tissue-level findings are concordant with the protein-level observations, linking cardiac-associated proteins to extracellular matrix remodelling and immune-related proteins to complement and inflammatory pathways. Consistent with these pathway-level observations, tissue and cell-type enrichment analysis demonstrated that upregulated proteins were enriched in fibroblast, myofibroblast, and smooth muscle cell populations, while downregulated proteins were enriched in plasma cell and immune-related compartments (Supplementary Fig. S7).

To assess whether these patterns were consistent across disease subtypes, we performed an exploratory clinician-guided stratification of amyloidosis cases based on linked clinical data and predefined subtype-specific criteria (see Methods and Supplementary Tables S3 and S4). This approach classified 31 cases as probable AL amyloidosis and 6 as probable ATTR amyloidosis, with the remainder unclassified. The proteomic signal was preserved in the AL subgroup, with consistent elevation of both cardiac-associated and immune-related proteins, whereas the ATTR subgroup showed greater variability, likely reflecting the limited sample size and precluding robust subtype-specific conclusions (Fig. 3b).

To further evaluate the specificity of the identified proteomic signature relative to a related plasma cell disorder, we compared protein expression between amyloidosis cases and individuals with multiple myeloma. After excluding individuals with both diagnoses (n = 10), the analysis included 165 myeloma and 51 amyloidosis cases. Several proteins, including FCRLB, MYBPC1, MYL3, NPPB, and NT-proBNP, remained significantly differentially expressed between the two groups following false discovery rate correction (Fig. 3c), whereas IGFBP1 and FABP1 were not.

Extending this comparison to a broader set of myeloma-associated proteins in the literature,^11^ revealed that canonical immune markers, including TNFRSF17, TNFRSF13B, SLAMF7, and TNFSF13B, were strongly enriched in the myeloma cohort, whereas cardiac-associated proteins remained more prominent in amyloidosis (Supplementary Fig. S3). These findings demonstrate both shared and disease-specific proteomic features, indicating that the observed signature is not solely reflective of plasma cell dyscrasia but includes distinct cardiac stress components characteristic of amyloidosis.

### Proteomic signatures are associated with time to diagnosis

To evaluate whether early plasma proteomic changes could predict the future diagnosis of amyloidosis at the individual level, we developed time-to-event prediction models integrating proteomic features with longitudinal outcome data. A schematic overview of the model development and evaluation workflow is shown in Fig. 4a. The dataset was randomly split into training (70%) and held-out test (30%) sets, with cases and controls split separately to avoid data leakage. All model development steps were performed exclusively using the training data.

**Figure 4.**
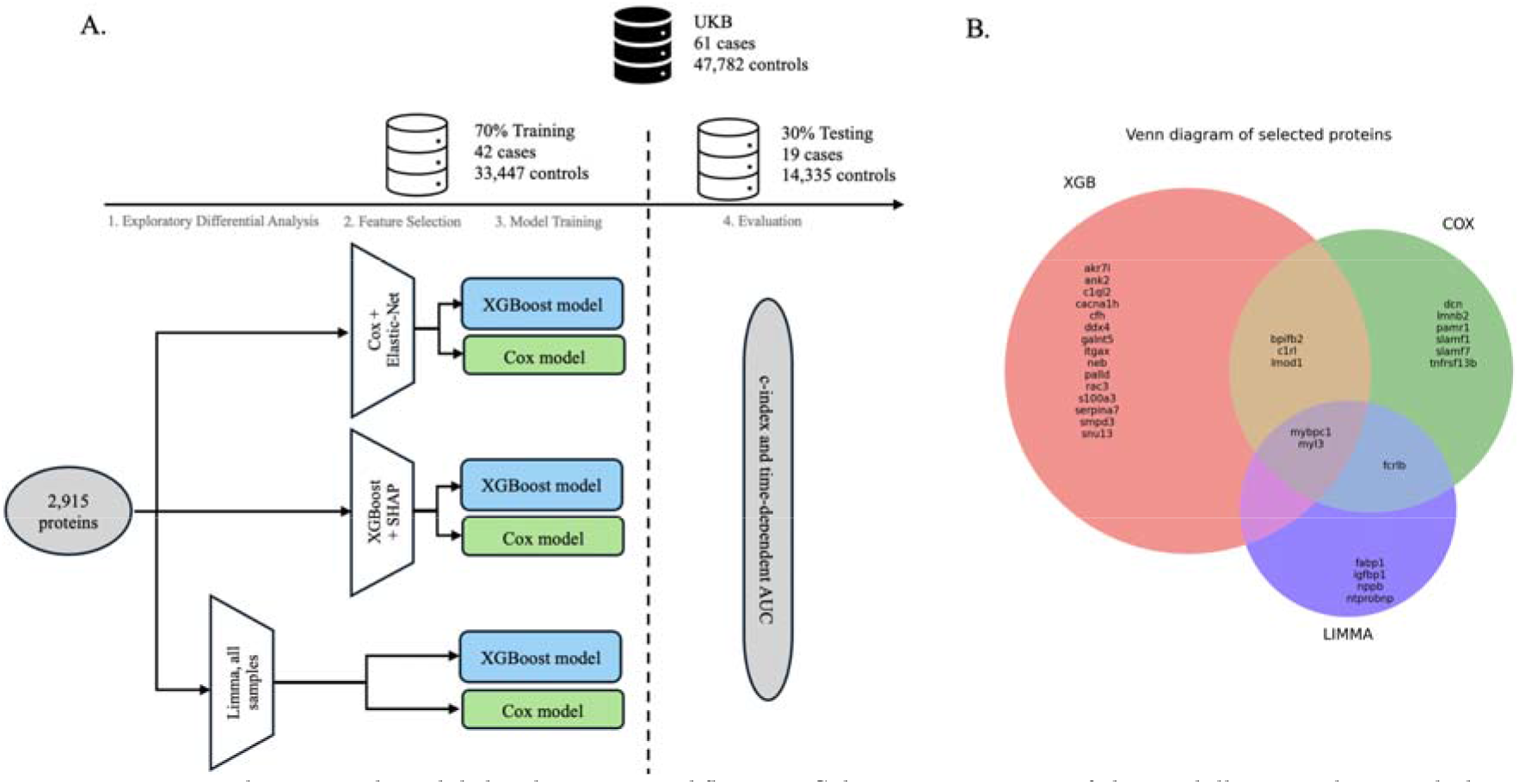
Feature selection and model development workflow. (a) Schematic overview of the modelling pipeline, including data partitioning, feature selection, model training, and evaluation. (b) Overlap of proteins identified by different feature selection methods, illustrating both shared and method-specific predictors.

As an initial step, we applied three complementary feature selection approaches to identify proteins associated with time to amyloidosis diagnosis: (i) univariate differential expression analysis (limma), (ii) elastic-net regularised Cox proportional hazards modelling, and (iii) gradient-boosted decision trees optimised for survival outcomes (XGBoost) with SHapley Additive exPlanations (SHAP) to quantify feature importance. Each approach identified partially overlapping sets of proteins, reflecting sensitivity to both linear and non-linear relationships (Fig. 4b).

To reduce redundancy and address multicollinearity, proteins with pairwise correlations greater than 0.9 were grouped, and representative features with higher variance were retained from each cluster during feature selection. This resulted in three distinct but overlapping sets of candidate proteins derived from each method.

For model development, we evaluated predictive performance using these method-specific feature sets. To further improve model stability, a more stringent correlation threshold (r > 0.7) was applied during final model training to derive parsimonious feature subsets.

The overlap between feature sets identified by different selection strategies is shown in Fig. 4b, highlighting both shared and method-specific predictors. Notably, cardiac-associated proteins such as MYL3 and MYBPC1 consistently emerged across methods, with additional overlap observed for immune-related proteins including FCRLB, supporting the presence of coherent biological signals underlying the predictive models.

### Robust prediction of amyloidosis risk up to 14 years before diagnosis

To quantify the temporal performance of proteomic risk prediction, we evaluated model discrimination across multiple time horizons using time-dependent area under the receiver operating characteristic curve (AUC) and concordance index (C-index) metrics on the held-out test set. Performance was assessed at 2-year intervals extending from early preclinical stages to near-diagnosis.

Both Cox proportional hazards and XGBoost survival models demonstrated sustained discriminative ability across extended follow-up periods (Fig. 5a). Time-dependent AUC values remained consistently high across all time horizons, indicating that predictive signal was not restricted to the period immediately preceding clinical diagnosis but was detectable many years in advance. Model performance remained stable across all time horizons with no evidence of substantial overfitting in the held-out test set.

**Figure 5.**
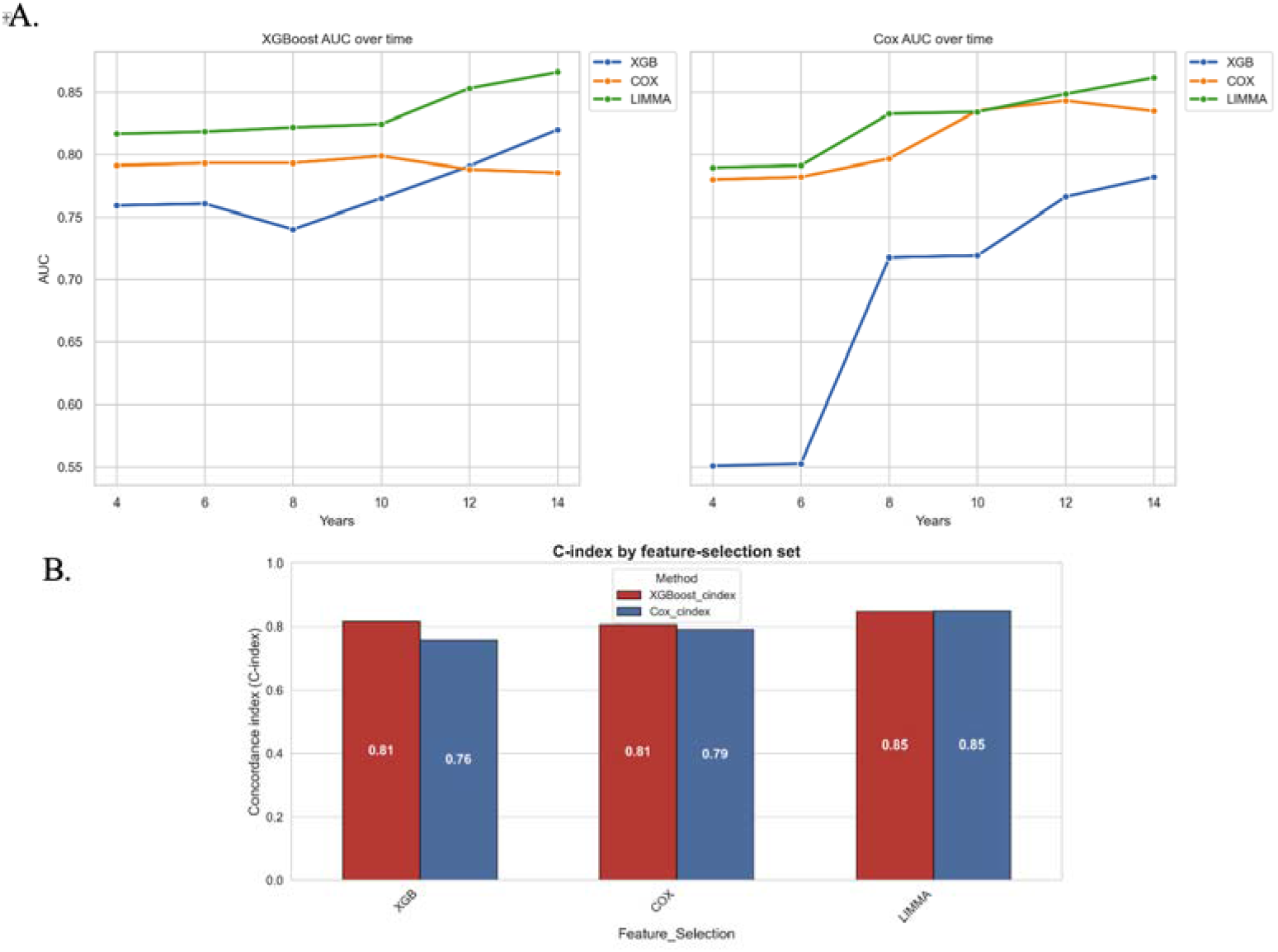
Robust prediction of amyloidosis risk across extended preclinical intervals. (a) Time-dependent area under the receiver operating characteristic curve (AUC) for amyloidosis prediction across multiple time horizons (4–14 years prior to diagnosis) using Cox proportional hazards and XGBoost survival models. Performance is shown for three feature selection strategies. (b) Concordance index (C-index) for models trained on different feature selection sets on the held-out test set.

Across feature selection strategies, models trained using proteins identified by differential expression analysis and refined through correlation filtering achieved the strongest and most consistent performance. Predictive performance was comparable across modelling approaches, with XGBoost showing modest improvements in some time horizons.

Concordance index comparisons in the held-out test set further supported the robustness of the predictive models (Fig. 5b). C-index values remained stable across feature selection strategies, with the highest performance observed for models based on differentially expressed proteins.

These findings indicate that amyloidosis risk can be predicted years before clinical diagnosis with stable performance across extended preclinical intervals, supporting the potential utility of plasma proteomics for longitudinal risk stratification.

Although the initial biological signature comprised seven proteins, predictive performance was largely preserved using a smaller subset of five proteins, indicating that the predictive signal is concentrated within a core group of biologically informative markers.

To assess the added predictive value of proteomic features, we compared models incorporating proteins with baseline models including only age and sex. Age- and sex-only models achieved moderate discrimination (C-index: 0.64 for XGBoost; 0.69 for Cox models), whereas inclusion of the parsimonious proteomic signature substantially improved performance (ΔC-index: 0.20 for XGBoost;

0.15 for Cox models). Similar improvements were observed for time-dependent AUC (ΔAUC: 0.15 for XGBoost; 0.08 for Cox models). These findings indicate that circulating protein markers provide substantial predictive information beyond demographic risk factors alone.

To interpret the final parsimonious model, we examined feature contributions using SHAP and estimated protein-specific effects using a Cox proportional hazards forest plot. SHAP analysis identified MYL3 as the strongest contributor to predicted risk, followed by age, FABP1, NPPB, FCRLB, and IGFBP1 (Fig. 6b), indicating that both cardiac and immune-related proteins contribute substantially to model predictions. The SHAP beeswarm plot further showed that higher values of key cardiac-associated proteins were generally associated with increased predicted risk (Fig. 6c), while the Cox model demonstrated a concordant direction of effect for the retained proteins (Fig. 6a). These results indicate that the parsimonious model preserves the internally consistent structure of the original proteomic signature.

**Figure 6.**
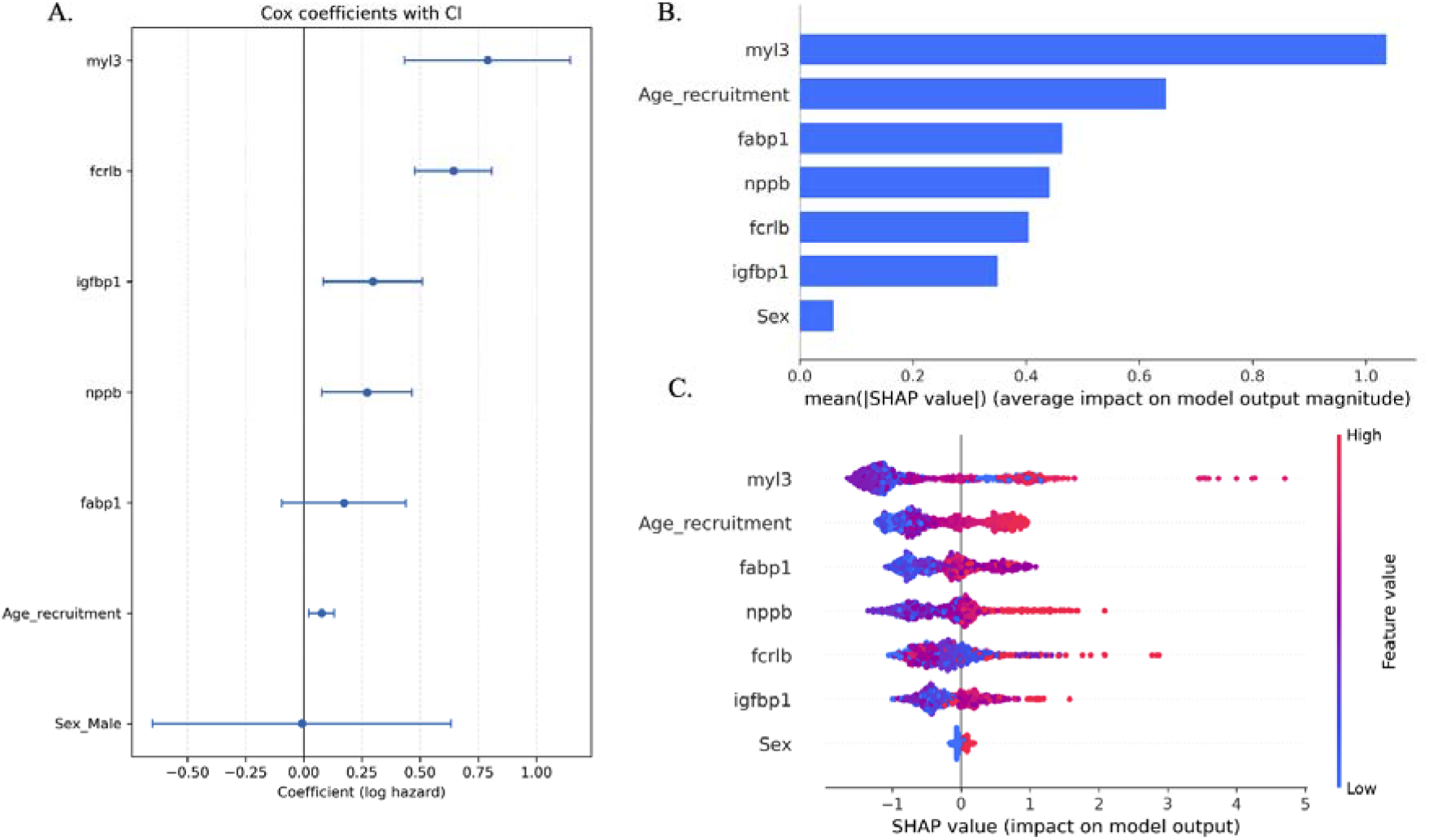
Interpretability of the final parsimonious prediction model. (a) Cox proportional hazards forest plot for the parsimonious protein set, showing effect sizes and 95% confidence intervals. (b) Mean absolute SHAP values for the selected proteins, showing their average contribution to model output. (c) SHAP beeswarm plot illustrating the direction and distribution of feature effects across individuals, with higher protein values generally associated with higher predicted risk for the strongest predictors.

### Exploratory analyses

We performed additional exploratory analyses to further assess the robustness and biological context of the identified proteomic signals.

Given emerging links between clonal haematopoiesis of indeterminate potential (CHIP), ageing, cardiovascular disease, and plasma cell disorders, we examined whether CHIP was enriched among amyloidosis cases within the subset of UK Biobank participants with available CHIP calls (n = 450,652). Among 420 amyloidosis cases with available CHIP data, 22 individuals harboured CHIP-associated mutations (∼5.24%), compared with 15,597 of 450,232 controls (∼3.46%). Despite this numerical difference, the relatively small number of CHIP-positive amyloidosis cases limited statistical power. In age- and sex-adjusted logistic regression models, neither CHIP presence nor CHIP burden was independently associated with incident amyloidosis. Variants in canonical CHIP-associated genes, including DNMT3A, TET2, and SRSF2, accounted for the majority of CHIP-positive cases, consistent with population-level distributions (Supplementary Fig. S10).

We also examined a small subset of individuals with prevalent amyloidosis at the time of protein assessment (n = 6) and compared their plasma proteomic profiles with those of individuals who developed amyloidosis during follow-up. No significant differences were observed between prevalent and incident cases for the key proteins identified in this study (Supplementary Fig. S11), suggesting that the detected proteomic signals are not confined to late-stage disease and are consistent across disease stages.

Together, these analyses support the robustness of the identified proteomic signature and indicate that it is not driven by clonal haematopoiesis or restricted to individuals with established disease at baseline.

### Proteomic signatures are not explained by pre-existing cardiac disease

Given the strong representation of cardiac-associated proteins among the early proteomic signals, we next assessed whether the observed associations could be explained by pre-existing cardiovascular disease rather than amyloidosis-specific processes.

Differential expression analysis adjusted for prior cardiac events demonstrated that several key proteins, including FCRLB, IGFBP1, MYL3, MYBPC1, and NPPB, remained significantly elevated in individuals who later developed amyloidosis (Fig. 7a). NT-proBNP also remained overexpressed, although with a modest reduction in effect size, whereas FABP1 was no longer significantly different. These findings indicate that the core proteomic signal is largely preserved after accounting for prior cardiac disease.

**Figure 7.**
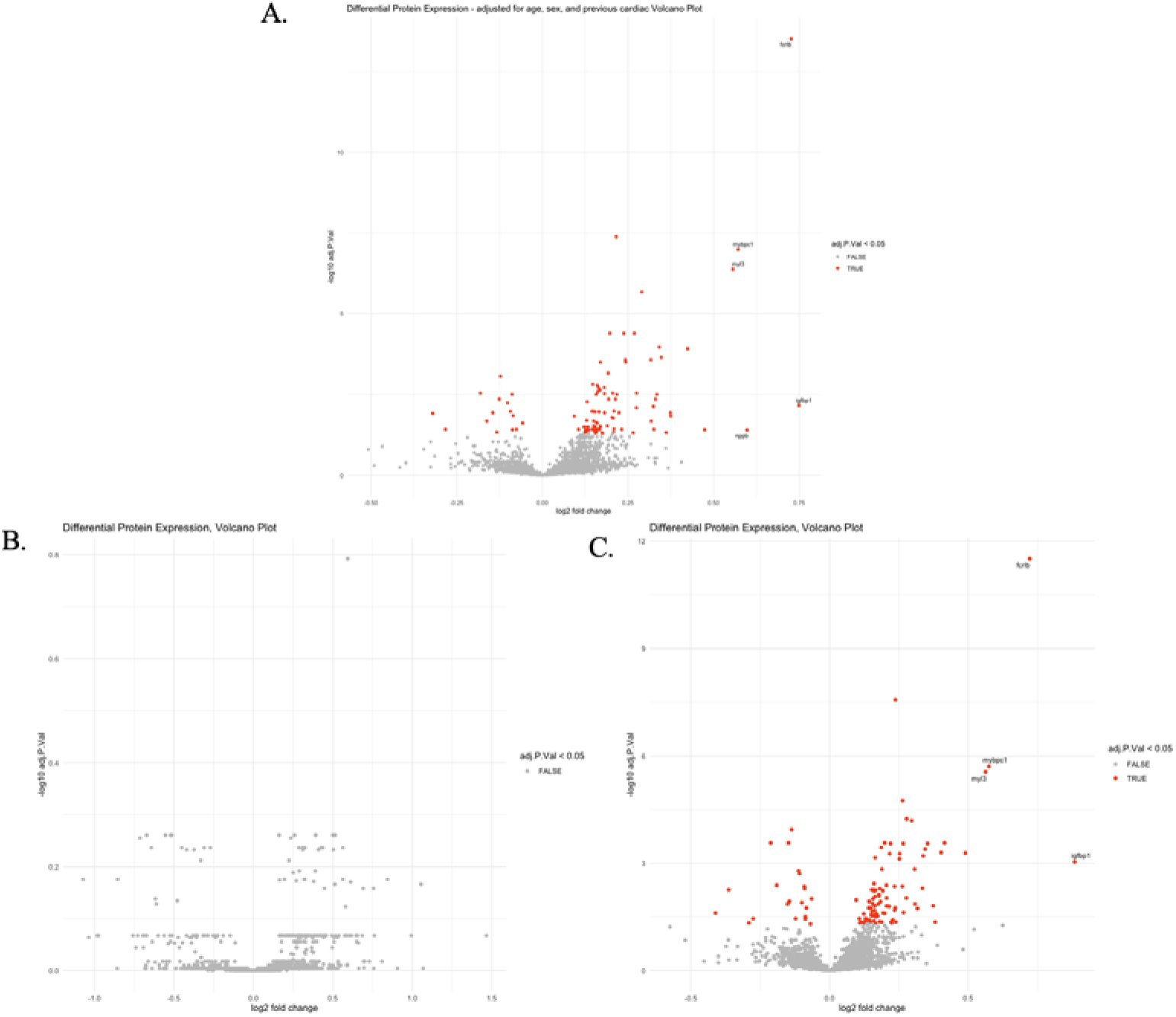
Proteomic signatures are not explained by pre-existing cardiac disease. (a) Differential protein expression analysis adjusted for age, sex, and prior cardiac events. (b) Comparison of plasma proteomic profiles in amyloidosis cases with and without prior cardiac events. (c) Comparison of selected proteins between amyloidosis cases and individuals with cardiac disease but no amyloidosis.

Among the 61 individuals who developed amyloidosis, 50 had at least one recorded cardiac event, occurring either before protein assessment (n = 15), between protein assessment and amyloidosis diagnosis (n = 30), or after diagnosis (n = 5) (Supplementary Fig. S4). We therefore compared plasma proteomic profiles between individuals with documented cardiac events prior to protein assessment and those without any recorded cardiac history. No significant differences in protein expression were observed between these groups (Fig. 7b), indicating that the early proteomic signals were not driven by pre-existing cardiac disease within the amyloidosis cohort.

We next compared amyloidosis cases to individuals with cardiac disease but no recorded amyloidosis. In this more stringent comparison, several proteins, including MYBPC1, MYL3, FCRLB, and IGFBP1, remained overexpressed in the amyloidosis cohort, whereas canonical cardiac biomarkers such as NT-proBNP and NPPB showed patterns consistent with broader cardiac pathology (Fig. 7c).

These analyses demonstrate that the observed proteomic alterations are not explained by pre-existing cardiovascular disease alone, but instead reflect disease-specific biological processes that precede clinically apparent amyloidosis.

## Discussion

### Principal findings

In this study, we demonstrate that systemic amyloidosis is preceded by plasma proteomic alterations detectable up to 14 years before clinical diagnosis. By combining population-scale proteomics with longitudinal follow-up data from the UK Biobank, we identified a set of seven circulating proteins that characterise the early biological signature of amyloidosis. From this set, a smaller parsimonious subset retained strong predictive performance for both the occurrence and timing of disease. These findings reveal related biological processes, including subclinical cardiac stress and immune dysregulation, that characterise the preclinical phase of disease.

Several key features define these findings. First, the temporal depth of prediction extends well beyond the timeframe in which current clinical symptoms become detectable. Second, the observed signals are not explained by overt cardiovascular disease or demographic confounding and remain distinct from those observed in related plasma cell disorders. Third, the proteomic patterns were internally consistent and reflected structured changes across biologically linked protein groups rather than stochastic variation.

Together, these results indicate that amyloidosis leaves a detectable molecular footprint in the circulation during a prolonged preclinical phase before clinical diagnosis, suggesting that disease-related biological processes are already active well before disease becomes clinically apparent.

### Biological interpretation

A central finding of this study is the elevation of cardiac-associated proteins several years before clinical diagnosis of amyloidosis. Proteins such as MYL3 and MYBPC1 were strongly co-expressed and consistently identified across both differential expression and predictive modelling approaches. In addition, natriuretic peptides, including NPPB and NT-proBNP, well-established markers of cardiac stress in overt amyloidosis, were elevated in individuals who were clinically unaffected at the time of sampling. Together, these findings suggest the presence of subtle, chronic myocardial stress prior to overt clinical recognition. These observations align with prior histopathological and imaging studies demonstrating that amyloid deposition can precede symptoms by many years and support a model in which early myocardial involvement is biologically active but clinically silent.

Importantly, cardiac proteins were not the sole drivers of the observed signal. Immune-related proteins, particularly FCRLB, were consistently elevated and showed weaker correlations with cardiac markers, suggesting a partially independent biological axis. FCRLB is primarily expressed in B cells and has been implicated in immune regulation and plasma cell biology, indicating that early immune dysregulation may contribute to amyloidosis risk. This is particularly relevant for AL amyloidosis, which arises from clonal plasma cell disorders.

Additional proteins, including IGFBP1 and FABP1, exhibited intermediate correlation patterns and were observed in both amyloidosis and myeloma cohorts, suggesting that they reflect broader systemic metabolic or inflammatory processes. Of note, these proteomic changes do not appear to reflect accelerated biological ageing, as no clear deviation from expected age-related proteomic trajectories was observed in heart or immune ageing models (Supplementary Fig. S9).

The parsimonious predictive models derived from these data retained representative proteins from both cardiac and immune components, indicating that the reduced model preserves the underlying biological structure of the full proteomic signature rather than reflecting a single dominant pathway. Additionally, these proteomic patterns were consistently observed in AL amyloidosis cases, suggesting that the signal is primarily driven by this subtype, while greater variability in ATTR cases likely reflects limited sample size, precluding robust subtype-specific conclusions.

Tissue and cell-type enrichment analysis of differentially expressed proteins further supported this interpretation, with upregulated proteins enriched in fibroblast and smooth muscle cell populations, consistent with extracellular matrix remodelling and early cardiac structural changes, while downregulated proteins were enriched in plasma cell and immune-related compartments, indicating interconnected contributions from cardiac structural remodelling and systemic immune processes.

Publicly available human heart single-cell atlases further support these findings, localising MYL3 and MYBPC1 to cardiomyocytes and NPPB to ventricular cardiomyocytes, while FCRLB shows minimal expression in cardiac tissue (Supplementary Fig. S8). These observations reinforce the interpretation that the observed plasma proteomic signature reflects linked contributions from cardiac structural and systemic immune compartments.

These findings are further supported by sensitivity analyses demonstrating that the observed proteomic alterations are not explained by pre-existing cardiac disease and are consistent across individuals with and without prior cardiac events. In addition, similar protein expression patterns were observed in individuals with prevalent amyloidosis at baseline compared with incident cases, indicating that these signals are not restricted to late-stage disease (Supplementary Fig. S11).

These findings support a model in which amyloidosis develops through interconnected biological pathways involving subclinical cardiac stress and immune dysregulation that are detectable long before clinical diagnosis.

### Comparison with prior work

Previous biomarker studies in amyloidosis have largely focused on patients at or near diagnosis, where biomarkers primarily reflect established organ involvement. Cardiac biomarkers such as NT-proBNP and troponins are invaluable for staging and prognosis but typically rise only after significant myocardial dysfunction has developed. In contrast, our findings demonstrate that cardiac-associated proteins are elevated many years earlier, at a stage when standard diagnostic pathways remain silent and individuals are clinically asymptomatic. This temporal distinction highlights the potential of these biomarkers for early disease detection.

Prior proteomic studies in amyloidosis have been predominantly cross-sectional, limited in scale, or focused on disease classification rather than prediction of incident disease. Such designs are inherently unable to distinguish early pathogenic signals from downstream consequences of amyloid deposition, organ dysfunction, or treatment. By leveraging longitudinal data from a large population cohort, the present study enables direct investigation of proteomic changes preceding clinical diagnosis, providing insight into early disease biology rather than late-stage pathology.

Our findings are consistent with observations in related plasma cell disorders, particularly multiple myeloma, in which dysregulation of immune and inflammatory proteins has been shown to precede clinical diagnosis by several years. These studies suggest that prolonged preclinical phases characterised by detectable molecular changes may represent a shared feature of protein misfolding and clonal plasma cell diseases. Extending this paradigm to amyloidosis represents an important conceptual advance.

### Clinical and translational implications

The identification of plasma proteomic signatures that predict future diagnosis of amyloidosis years before clinical presentation has important clinical and translational implications. Early identification of individuals at increased risk could enable targeted surveillance, earlier diagnostic evaluation, and intervention at a stage when organ damage may still be preventable. This is particularly relevant for high-risk populations, including individuals with monoclonal gammopathy of undetermined significance (MGUS) and patients with heart failure with preserved ejection fraction (HFpEF), in whom amyloidosis is often under-recognised.

Notably, predictive performance was maintained using a parsimonious subset of proteins, suggesting that early risk stratification may not require large-scale proteomic panels. Instead, a limited number of mechanistically linked markers may capture much of the predictive signal. However, translation into clinical practice will require external validation in independent cohorts, as well as calibration of thresholds using platforms that provide absolute protein quantification.

More broadly, these findings support a shift from reactive diagnosis to proactive risk assessment in amyloidosis. Rather than relying on the development of overt organ dysfunction, population-level molecular profiling may enable identification of disease risk during a prolonged preclinical phase, potentially allowing earlier intervention and improved patient outcomes.

### Future directions

Future work should focus on validating these findings in independent clinical and population-based cohorts with proteomic data and long-term follow-up. In particular, validation in well-characterised amyloidosis cohorts will be essential to confirm clinical relevance for disease progression, subtype specificity, and generalisability of the identified biomarkers. Translation of these findings into clinical practice will require validation using platforms that enable absolute protein quantification.

Further studies integrating proteomic data with genomic and transcriptomic analyses may help clarify the biological pathways underlying early disease development. In particular, tissue-level and single-cell approaches could provide insight into the cellular origins of the observed proteomic signals and the interplay between subclinical cardiac stress and immune dysregulation in the preclinical phase of amyloidosis.

Prospective studies in high-risk populations, including individuals with monoclonal gammopathy of undetermined significance (MGUS) and patients with heart failure with preserved ejection fraction, will be critical to evaluate the clinical utility of early proteomic risk stratification and to define its role in guiding screening and early intervention strategies.

## Methods

### Study population, data sources, and plasma proteomic profiling

This study was conducted using data from the UK Biobank, a large prospective population-based cohort comprising over 500,000 individuals aged 40–69 years at recruitment. Participants provided written informed consent, and ethical approval was obtained by the UK Biobank from the relevant ethics committees. The present analyses were conducted under an approved UK Biobank application.

Plasma proteomic data were available for a subset of participants as part of the UK Biobank Pharma Proteomics Project (UKB-PPP), a large-scale proteomics initiative. Proteomic measurements were linked to longitudinal health outcomes through hospital episode statistics, primary care records, cancer registry data, and death registry data. Demographic variables, including age and sex, were obtained from baseline assessment data.

Plasma protein measurements were generated using a high-throughput affinity-based proteomics platform (Olink Proteomics), which utilises proximity extension assay technology coupled with next-generation sequencing for protein quantification. Protein expression levels were reported as normalised protein expression (NPX) values on a log_2_ scale. Quality control and normalisation were performed according to UKB-PPP protocols. Proteins failing quality control or with low detection rates were excluded, resulting in approximately 2,900 proteins per participant. Missing values were imputed using median imputation within case and control groups for differential expression analyses.

### Definition of the disease and index date

Amyloidosis cases were identified using linked electronic health records and defined using International Classification of Diseases, Tenth Revision (ICD-10) codes corresponding to systemic amyloidosis (E85.4, E85.8, and E85.9). Participants with a recorded diagnosis of amyloidosis prior to plasma protein assessment (n = 6) were excluded to ensure inclusion of incident cases only.

The date of plasma protein assessment was defined as the index date for all analyses. Time to diagnosis was calculated as the interval between the index date and the first recorded amyloidosis diagnosis. Participants without a recorded diagnosis of amyloidosis in linked health records or death registry data were censored at the end of available follow-up.

### Differential protein expression analysis

Differential protein expression between individuals who later developed amyloidosis and controls was assessed using linear models with empirical Bayes moderation (limma). Analyses were performed both unadjusted and adjusted for age and sex at the index date. Log_2_ fold changes (LFC), moderated t-statistics, p-values, and false discovery rate (FDR)–adjusted p-values were calculated. Proteins with LFC > 0.5 and FDR-adjusted p-values < 0.05 were considered differentially expressed.

### Unsupervised analyses and correlation structure

Principal component analysis (PCA) was performed on scaled proteomic data to explore global patterns of variation and assess overall structure in the dataset. Pairwise correlations among selected proteins were calculated using Pearson correlation coefficients and visualised using correlation matrices.

### Time-to-event predictive modelling

To evaluate the ability of plasma proteomic signatures to predict future amyloidosis, time-to-event models were developed using Cox proportional hazards models and gradient-boosted survival trees (XGBoost). XGBoost models were trained using a Cox proportional hazards objective function. Cox models were used for interpretable hazard estimation, while XGBoost models were used to capture non-linear relationships and interactions between proteins.

Given the low number of incident events, model complexity was controlled through feature selection, regularisation, and validation on a held-out test set. Model complexity was further constrained through shallow tree depth in XGBoost and regularisation in Cox models.

Prior to data partitioning, controls younger than the youngest amyloidosis case minus two years were excluded to ensure comparability of age distributions between cases and controls and to avoid extrapolation beyond the observed case age range.

The dataset was partitioned once into a training set (70%) and a held-out test set (30%). Cases and controls were split separately to prevent data leakage, ensuring that individuals contributing to model development were not present in the evaluation set. The held-out test set was not used in any feature selection, model development, or resampling procedures.

Within the training set, model development was performed using cross-validation and bootstrap resampling. To mitigate class imbalance, each case was matched to controls (1:5 ratio) based on age and sex within bootstrap samples. These matched datasets were used to estimate feature stability.

Cross-validation was used for model evaluation within the training set, while bootstrap resampling was used independently to assess feature stability; no additional train–test splits were performed within bootstrap iterations.

### Feature selection

Feature selection was performed using complementary approaches. For XGBoost models, features were ranked by mean absolute SHAP values across cross-validation folds, and the top 20 features were retained. For Cox proportional hazards models, feature stability was assessed across bootstrap resamples, and features were retained if they were selected in at least 70% of bootstrap iterations.

These strategies ensured that selected features reflected both predictive importance and stability across resampling procedures.

### Multicollinearity control

To reduce redundancy, proteins with absolute pairwise correlation >0.9 were grouped, and the highest-variance representative from each cluster was retained. For final model construction, an additional correlation threshold (|r|>0.7) was applied to derive parsimonious feature sets.

### Final model training

After feature selection and tuning, final models were refit on the entire training dataset using the selected feature sets. XGBoost survival models were trained using the optimised hyperparameters and selected features. Cox proportional hazards models were refit using the same selected features. These final models were then evaluated once on the held-out test set.

### Incremental predictive value

To quantify the contribution of proteomic features beyond demographic risk factors, baseline models including only age and sex were compared to models incorporating proteomic features using differences in C-index and time-dependent AUC.

### Model interpretability

Model interpretability for XGBoost models was assessed using SHapley Additive exPlanations (SHAP). SHAP values were used in two complementary ways. First, feature importance was quantified across cross-validation folds using mean absolute SHAP values, and these rankings were used for feature selection. Second, SHAP values were computed for the final parsimonious model to characterise the contribution, direction, and distribution of each feature’s effect on predicted risk at the individual level.

For Cox proportional hazards models, predictors were standardised prior to model fitting to improve numerical stability. Regression coefficients were estimated on the standardised scale and subsequently back-transformed to hazard ratios per unit increase in the original variables. These transformed estimates were used for forest plot visualisation.

### Model evaluation

Model performance was evaluated exclusively on the held-out test set. The concordance index (C-index) and the time-dependent area under the receiver operating characteristic curve (AUC) were assessed independently. AUC was evaluated at prespecified horizons ranging from 4 to 14 years at 2-year intervals.

Comparisons were made between baseline (age and sex) and clinical-plus-proteomic models to assess the added predictive value of proteomic features.

### Subtype classification of amyloidosis cases

Amyloidosis cases were stratified into probable AL and ATTR subtypes using clinician-guided annotation based on linked health records. Classification criteria included diagnostic codes, procedures, and associated clinical conditions (Supplementary Tables S3–S4). Cases not meeting criteria were classified as unassigned.

### Cardiac sensitivity

To assess whether observed proteomic associations were driven by pre-existing cardiovascular disease, sensitivity analyses were performed adjusting for documented cardiac events prior to the index date. Additional analyses stratified amyloidosis cases by cardiac involvement and compared amyloidosis-associated proteomic signatures with those observed in individuals with cardiac disease but no recorded amyloidosis.

### CHIP analysis

Clonal haematopoiesis of indeterminate potential (CHIP) was assessed in the subset of participants with available CHIP calls derived from genomic sequencing data. CHIP was defined based on the presence of somatic variants in established CHIP-associated genes above predefined variant allele frequency thresholds. Logistic regression models adjusted for age and sex were used to evaluate associations between CHIP status, CHIP burden, and incident amyloidosis. Given the limited number of CHIP-positive amyloidosis cases, these analyses were considered exploratory and not intended to support causal inference.

## Data Availability

UK Biobank data are available to bona fide researchers upon application to the UK Biobank. Proteomic data were accessed through approved UK Biobank resources.

## Code Availability

Custom code used for data processing, statistical analysis, and model development is available from the corresponding author upon reasonable request.

## References

1. Wechalekar, A. D., Gillmore, J. D. & Hawkins, P. N. Systemic amyloidosis. Lancet 387, 2641–2654 (2016).

2. Garcia-Pavia, P. et al. Diagnosis and treatment of cardiac amyloidosis: a position statement of the ESC Working Group on Myocardial and Pericardial Diseases. Eur. Heart J. 42, 1554–1568 (2021).

3. Dispenzieri, A. et al. Serum cardiac troponins and N-terminal pro–brain natriuretic peptide: a staging system for primary systemic amyloidosis. J. Clin. Oncol. 22, 3751–3757 (2004).

4. Hahn, V. S. et al. Endomyocardial biopsy characterization of heart failure with preserved ejection fraction and prevalence of cardiac amyloidosis. JACC Heart Fail. 8, 712–724 (2020).

5. Mangiacavalli, S. et al. Feasibility of a biomarker-based screening for pre-symptomatic AL amyloidosis in patients with intermediate/high-risk MGUS. Am. J. Hematol. 100, 342–345 (2025).

6. Ibrahim Abdalla Ibrahim, F. et al. The role of artificial intelligence in the detection of cardiac amyloidosis: a systematic review. Cureus 17, e78488 (2025).

7. Ahmadi-Hadad, A. et al. Artificial intelligence as a tool for diagnosis of cardiac amyloidosis: a systematic review. J. Med. Biol. Eng. 44, 499–513 (2024).

8. Grogan, M. et al. Value of artificial intelligence for enhancing suspicion of cardiac amyloidosis using electrocardiography and echocardiography: a narrative review. J. Am. Heart Assoc. 14, e036533 (2025).

9. Allegra, A. et al. Machine learning approaches in diagnosis, prognosis and treatment selection of cardiac amyloidosis. Int. J. Mol. Sci. 24, 5680 (2023).

10. Royer, P. et al. Large-scale plasma proteomics in the UK Biobank modestly improves prediction of major cardiovascular events in a population without previous cardiovascular disease. Eur. J. Prev. Cardiol. 31, 1681–1689 (2024).

11. Fieggen, J. et al. Dysregulated immune proteins in plasma in the UK Biobank predict multiple myeloma 12 years before clinical diagnosis. Blood Adv. 9, 3766–3770 (2025).

