## Supplementary Materials for "Early plasma proteomic alterations precede amyloidosis diagnosis, reflecting cardiac and immune dysregulation"

Table S1. Baseline characteristics of the full proteomics cohort, comparing controls and amyloidosis cases.

|  | | Controls | Amyloidosis | p-value |
| --- | --- | --- | --- | --- |
| N | | 47782 | 61 |  |
| Age at recruitment (mean (SD)) | | 57.20 (8.16) | 62.75 (6.04) | <0.001 |
| Sex (%) | **Female** | 25909 (54.2) | 29 (47.5) | 0.358 |
|  | **Male** | 21873 (45.8) | 32 (52.5) |  |

Table S2. Baseline characteristics of the proteomics subset with available proteomic data, showing representativeness relative to the full cohort.

|  | | Amyloidosis Without Protein | Amyloidosis With Protein | p-value |
| --- | --- | --- | --- | --- |
| N | | 422 | 61 |  |
| Age at recruitment (mean (SD)) | | 62.37 (6.16) | 62.75 (6.04) | 0.65 |
| Sex (%) | **Female** | 184 (43.6) | 0.659 | 0.358 |
|  | **Male** | 238 (56.4) | 32 (52.5) |  |

Table S3. Diagnostic and procedural codes used to define probable AL amyloidosis by clinician-guided annotation.

| **Disease / Condition** | **ICD-10** | **ICD-9** | **OPCS-3** | **OPCS-4** |
| --- | --- | --- | --- | --- |
| **Multiple myeloma** | C90.0 | 203 | – | – |
| **Monoclonal gammopathy** | D47.2 | 273.1 |  |  |
| **Non-Hodgkin lymphoma (various subtypes)** | C82,C83,C84, C85, C85.1, C85.2, C85.8, C85.9,C86,C88,C91,D47 | 200.8, 200.9 | – | – |
| **Waldenström macroglobulinemia** | C88.0 | 202.4 | – | – |
| **Plasma cell leukemia** | C90.1 | 203.1 | – | – |
| **Disorder involving the immune mechanism, unspecified** | D89.9 |  |  |  |
| **Bone marrow biopsy** | – | – | 798.2 | Y20 |
| **Bone marrow transplant** | Z94.81 *(not searchable)* | V42.81 | 798.3 | W35.8 |
|  |  |  |  | Y69.8 |
| **Autologous stem cell transplant** | – | – | – | X33.4 |
| **Chemotherapy (encounter/delivery)** | Z51.1 *(not searchable)* | V58.11, V58.12 | 961.1 | X72–X74, X74, X70–X74, X29.2 |
| **Long-term immunosuppressant use** | Z79.6 *(not searchable)* | V58.69 | – | – |
|  | *D70* |  |  |  |
| **PET-CT** | – | – | – | U36.2 |
| **Nephrotic syndrome** | N04 | 581 | – |  |
| **Macroglossia** | Q38.2 , C02.9 | 750.15 | – | – |
| ***Periorbital purpura*** | S00.1 | 920 | – | – |
| ***Morbidity -Mortality code (Complications of medical and surgical care)for antineoplastic systemic agent*** | Y43.1,Y43.2,Y43.3,Y43.4,Y43.5,Y43.8,Y43.9 |  |  |  |
| ***abnormalities of plasma proteins*** | *R70.1, R77.0, R77.1 ,R77.8, R77.9* | 790.6 |  |  |

Table S4. Diagnostic and procedural codes used to define probable ATTR amyloidosis by clinician-guided annotation.

| **Disease / Condition** | **ICD-10** | **ICD-9** | **OPCS-3** | **OPCS-4** |
| --- | --- | --- | --- | --- |
| **Carpal tunnel syndrome** | G56.0 | 354 | 41.1 | W76.1 |
| **Spinal stenosis** | M48.0 | 724.01–724.03 | 020.5, 022 | V25.3, V25.4, V25.5, V29.6, V33.7 |
| **Tendon rupture (non-traumatic)** | M66 | 727.60–727.63 | 842,845.3, 846.3, 848.3, 848, 852, 852.3 | T67.5, T67.6, T68.2 |
| **Visual loss (unspecified)** | H54.7 | 369.9 | – | – |
| **Spinal deformity / stabilization surgery** | – | – | – | V41, V40 |
| **Tendon repair/reconstruction (hand)** | – | – | 846.3, 848.3, 852.3 | T67.5, T67.6, T68.2 |

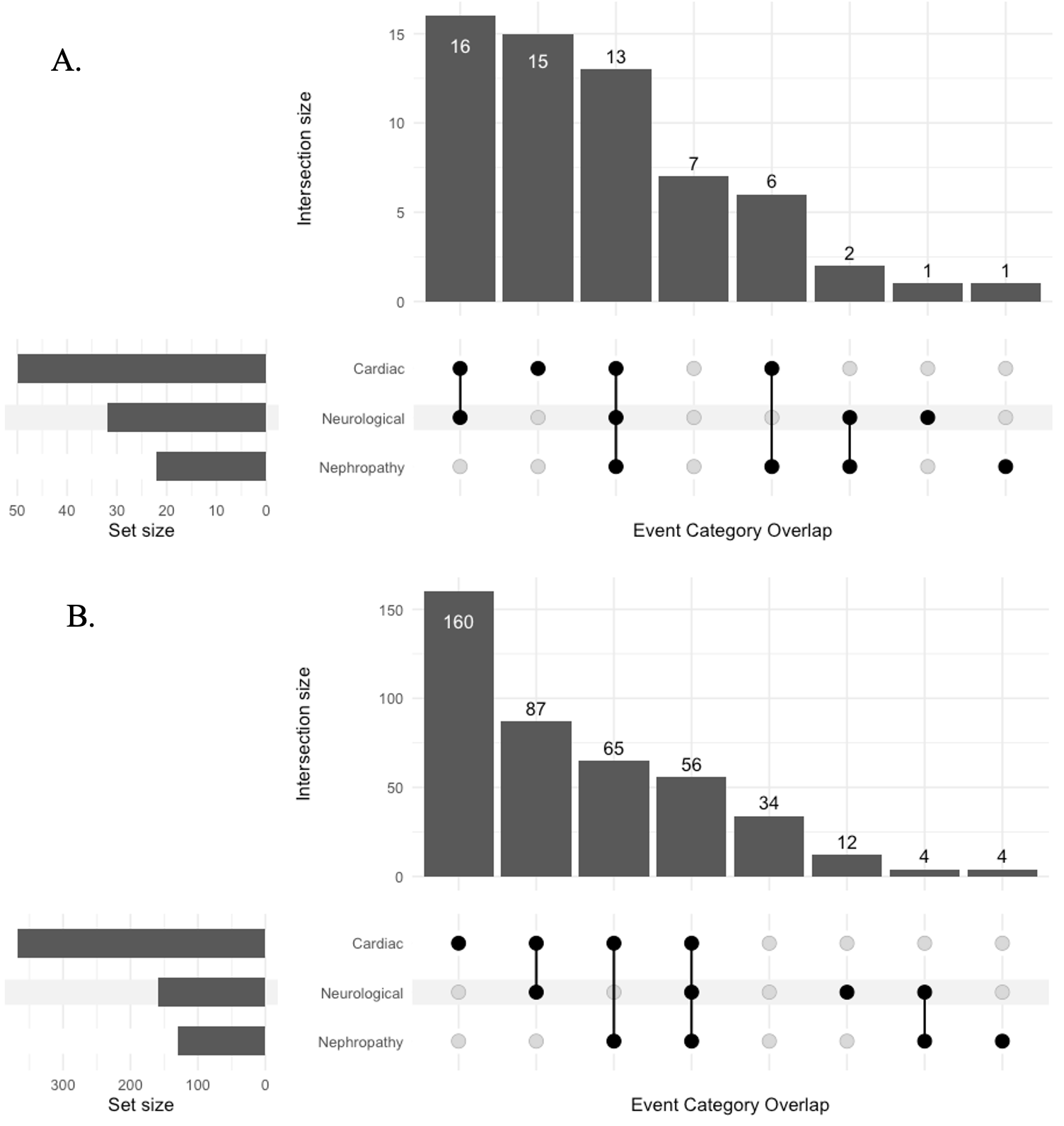

Figure S1. Comparison of amyloidosis-related comorbidity patterns in participants with and without proteomic data. a. Event overlap structure in amyloidosis cases with proteomic data. b. Event overlap structure in participants without proteomic data.

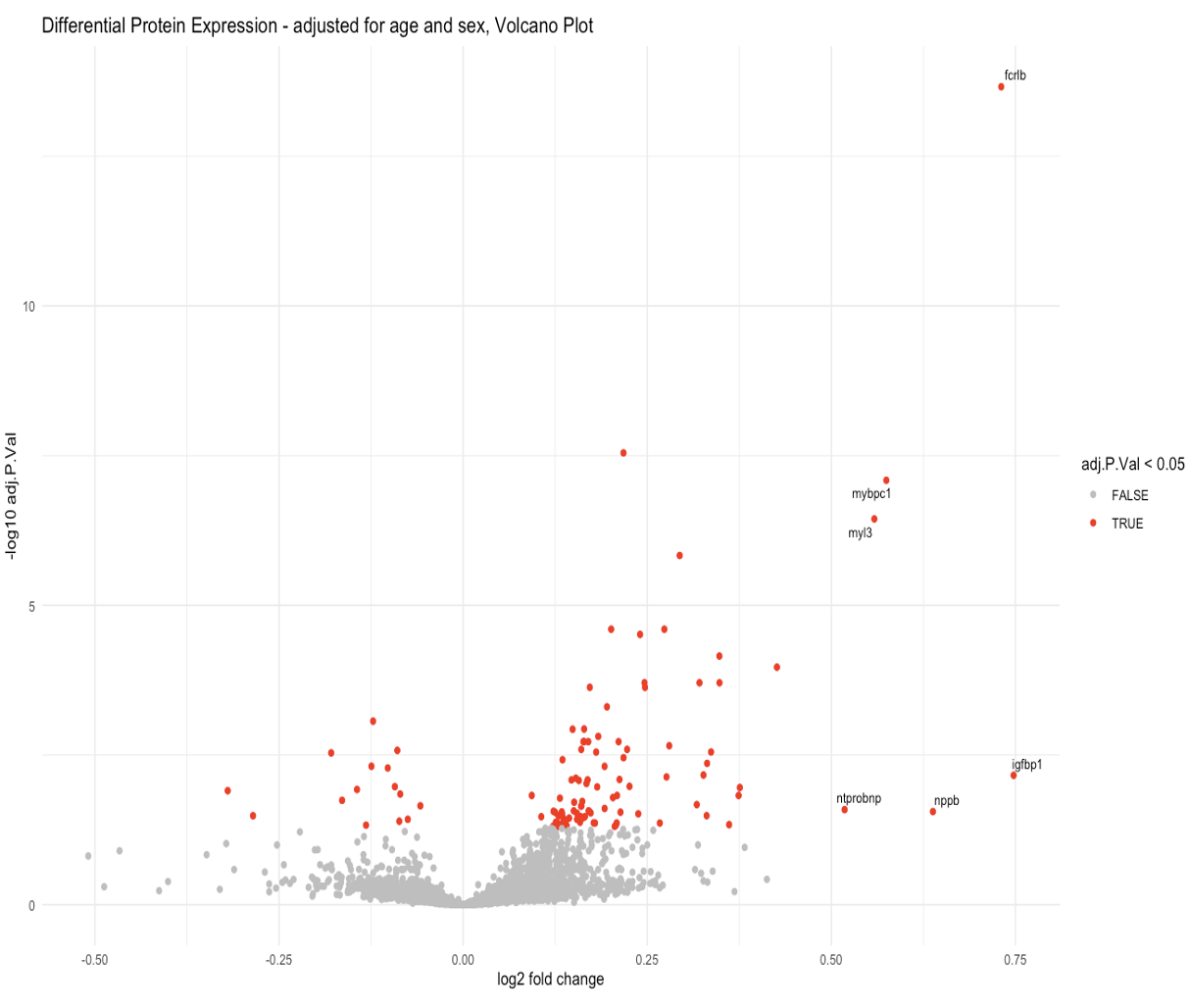

*Figure S2. Age- and sex-adjusted differential protein expression analysis comparing amyloidosis cases with controls. Significantly associated proteins are highlighted.*

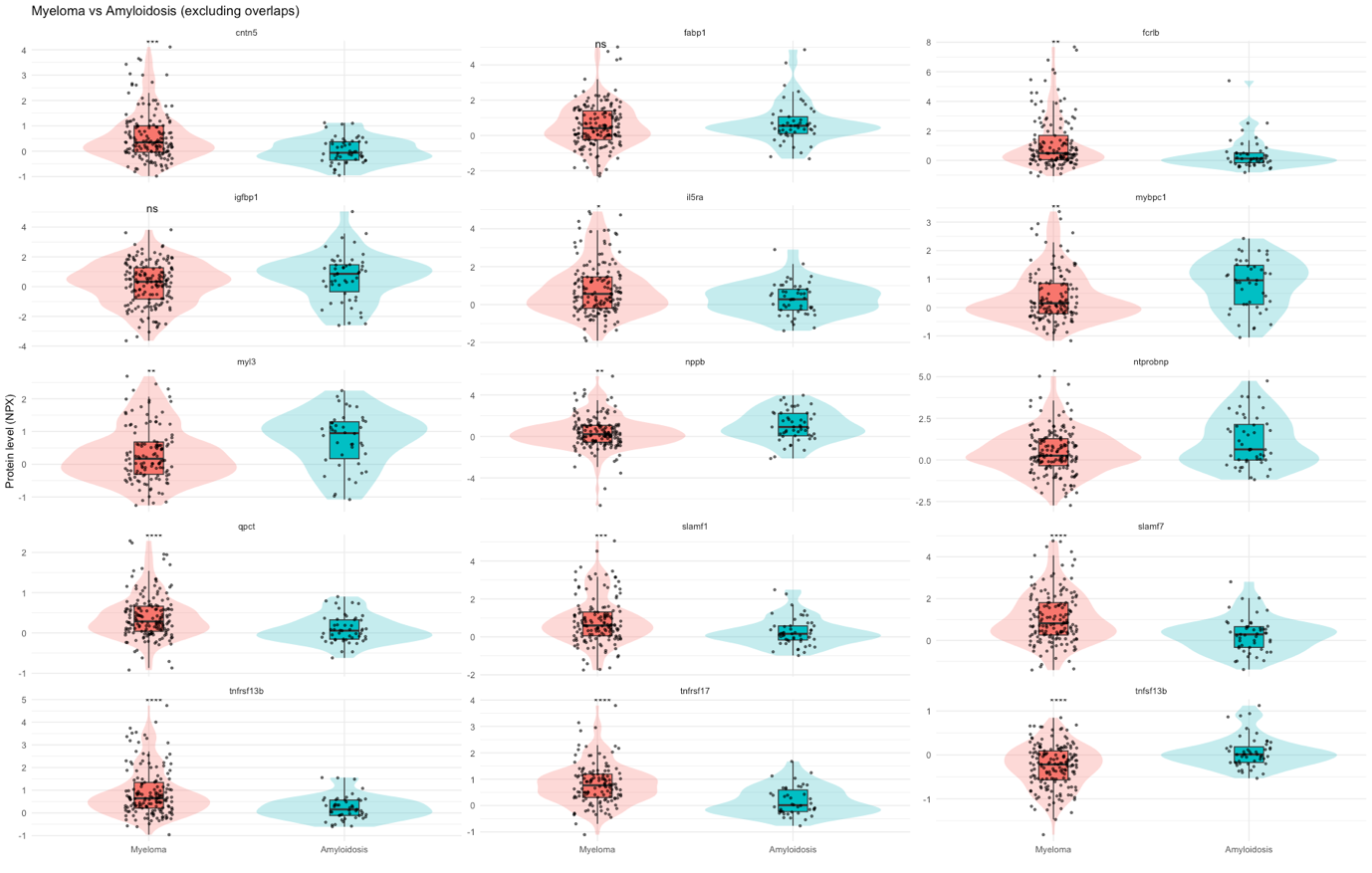

*Figure S3. Comparison of amyloidosis- and myeloma-associated protein expression, highlighting shared and disease-specific signals. In each box, the distribution of selected proteins across amyloidosis and myeloma groups.*

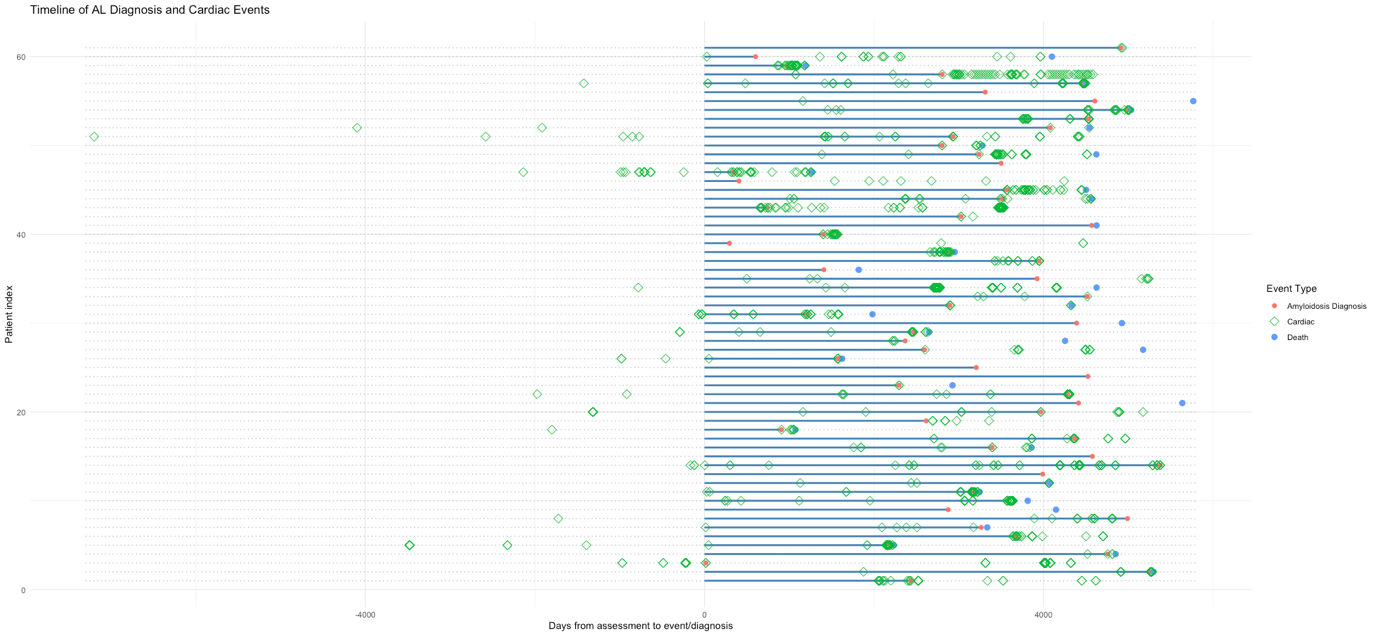

Figure S4. Timing of amyloidosis diagnosis, cardiac events, and death among proteomics participants, providing the clinical context for the longitudinal analyses.

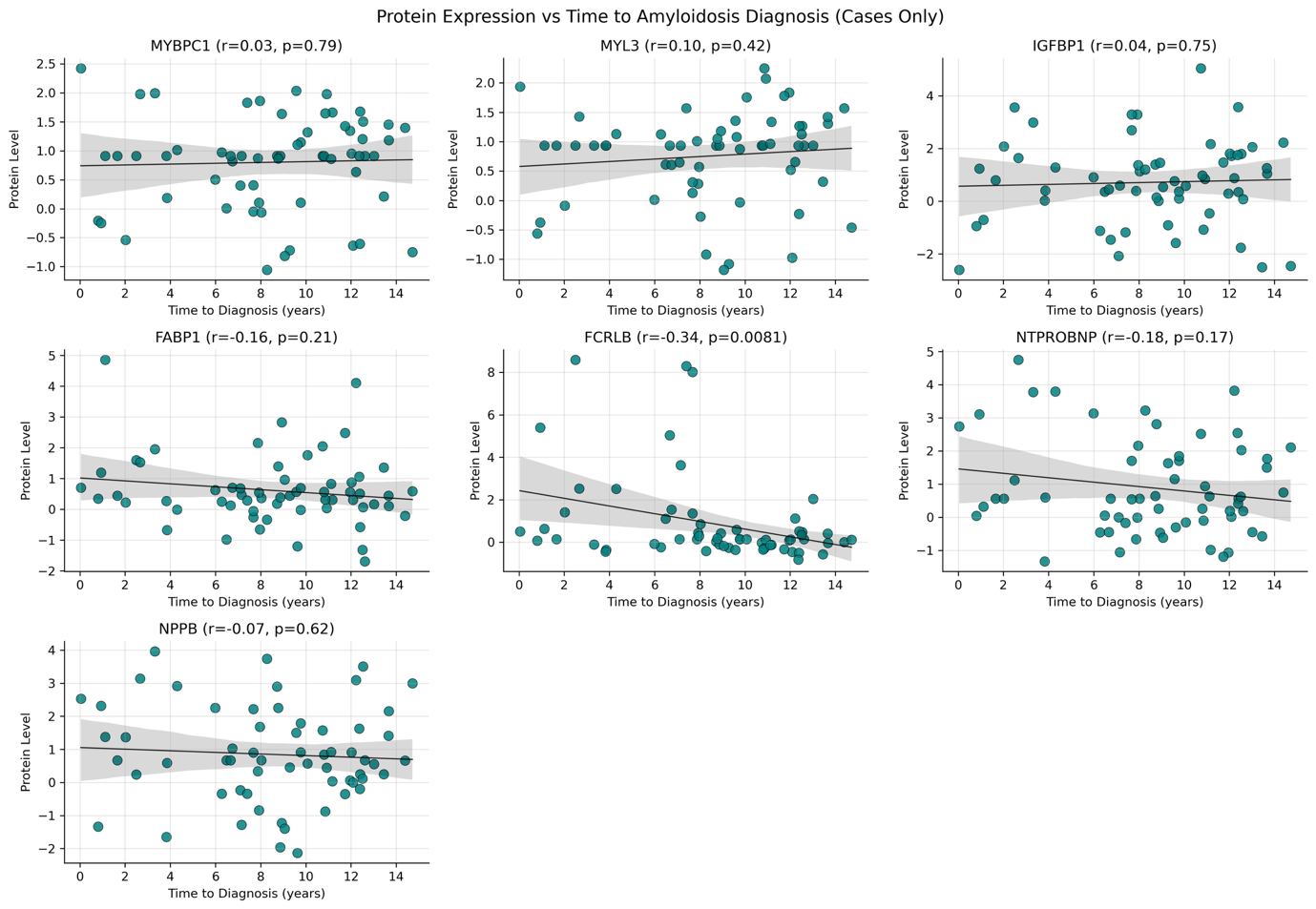

Figure S5. Longitudinal trajectories of key proteins relative to time to amyloidosis diagnosis, showing progressive protein elevation before clinical recognition.

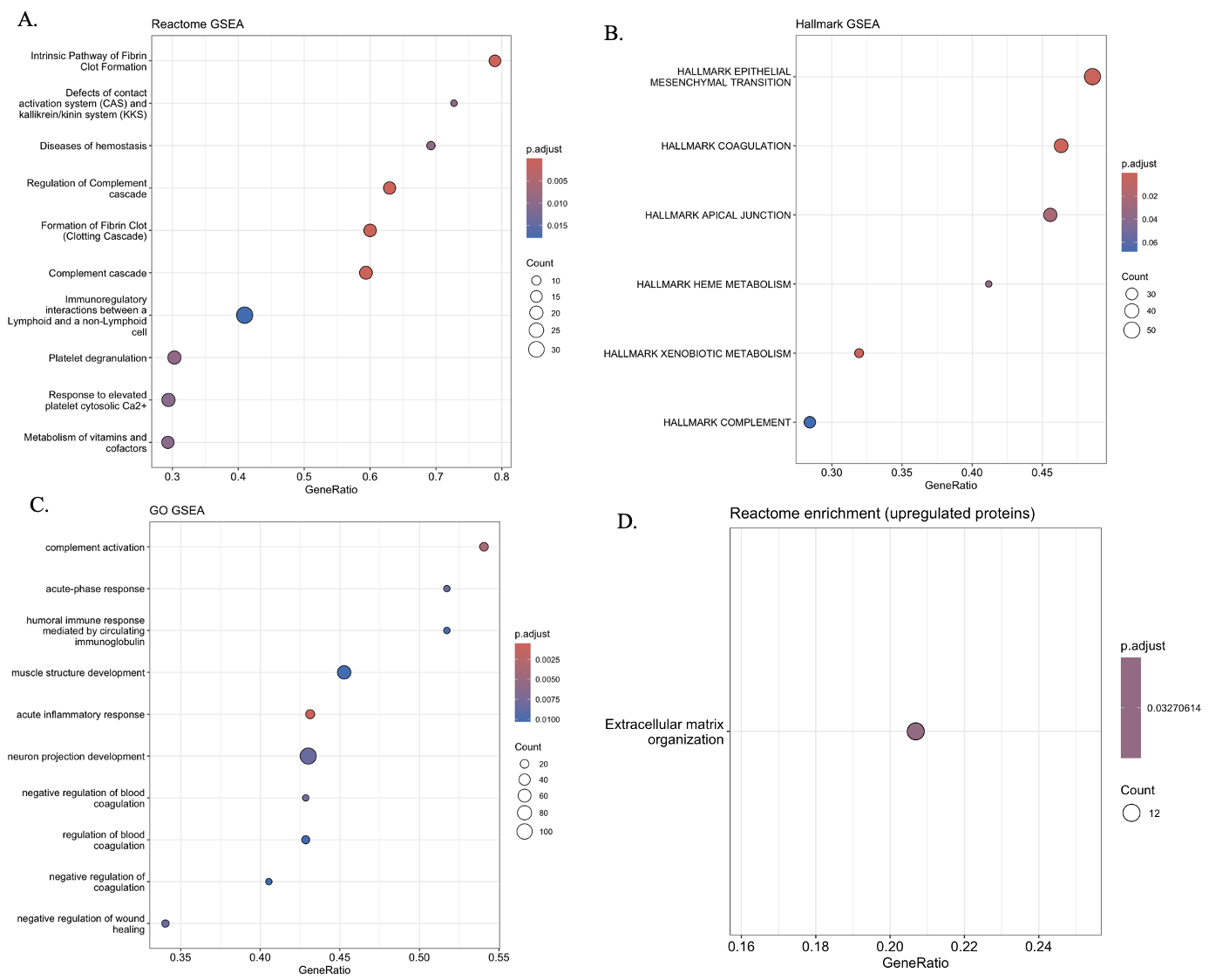

Figure S6. Pathway enrichment analysis integrating ranked (GSEA) and over-representation (ORA) approaches, highlighting extracellular matrix remodelling, cardiac development, coagulation, complement, and inflammatory pathways. a. Reactome GSEA results (rank-based enrichment across the full proteome). b. Hallmark GSEA results summarising major biological programmes. c. Gene Ontology (GO BP) GSEA results. d. Reactome over-representation analysis (ORA) of upregulated proteins.

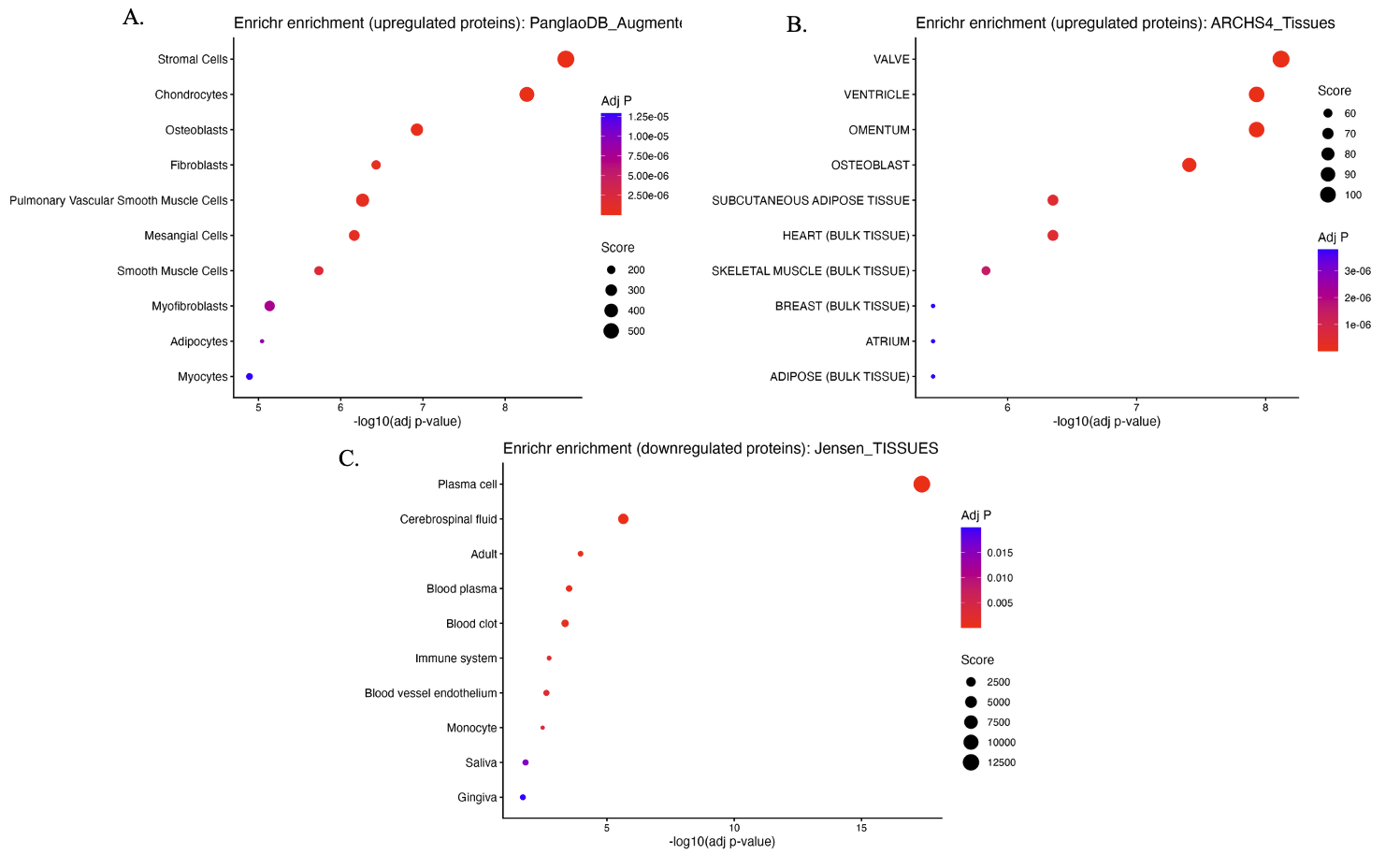

Figure S7. Tissue and cell-type enrichment analysis of differentially expressed proteins using multiple reference datasets. a. PanglaoDB cell-type enrichment of upregulated proteins. b. ARCHS4 tissue enrichment of upregulated proteins. c. Jensen Tissues enrichment of downregulated proteins.

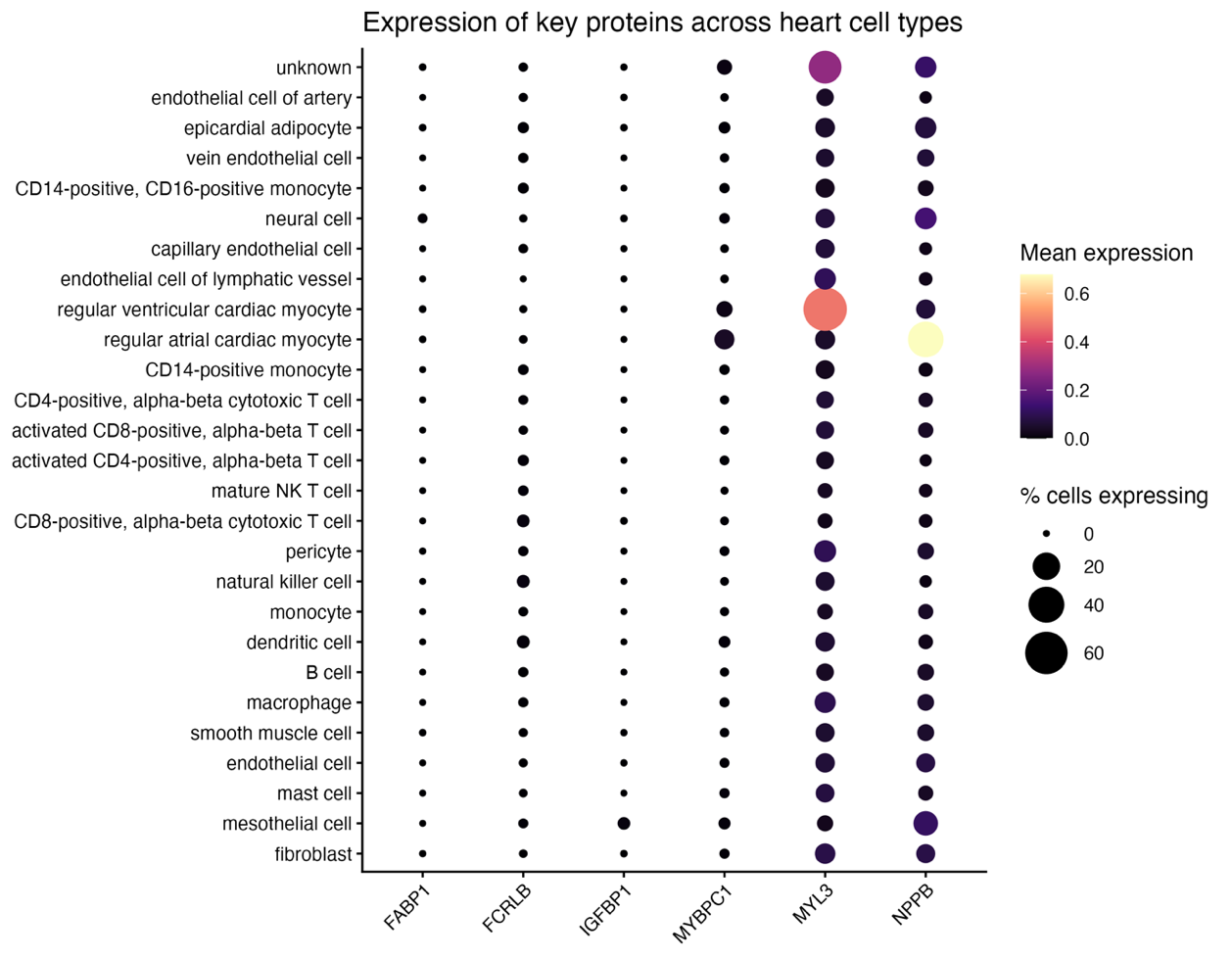

Figure S8. Expression of key proteins across human heart cell types in a public single-cell atlas. MYL3, MYBPC1, and NPPB show cardiomyocyte-specific expression, whereas FCRLB shows minimal expression in cardiac cell populations, supporting its non-cardiac origin.

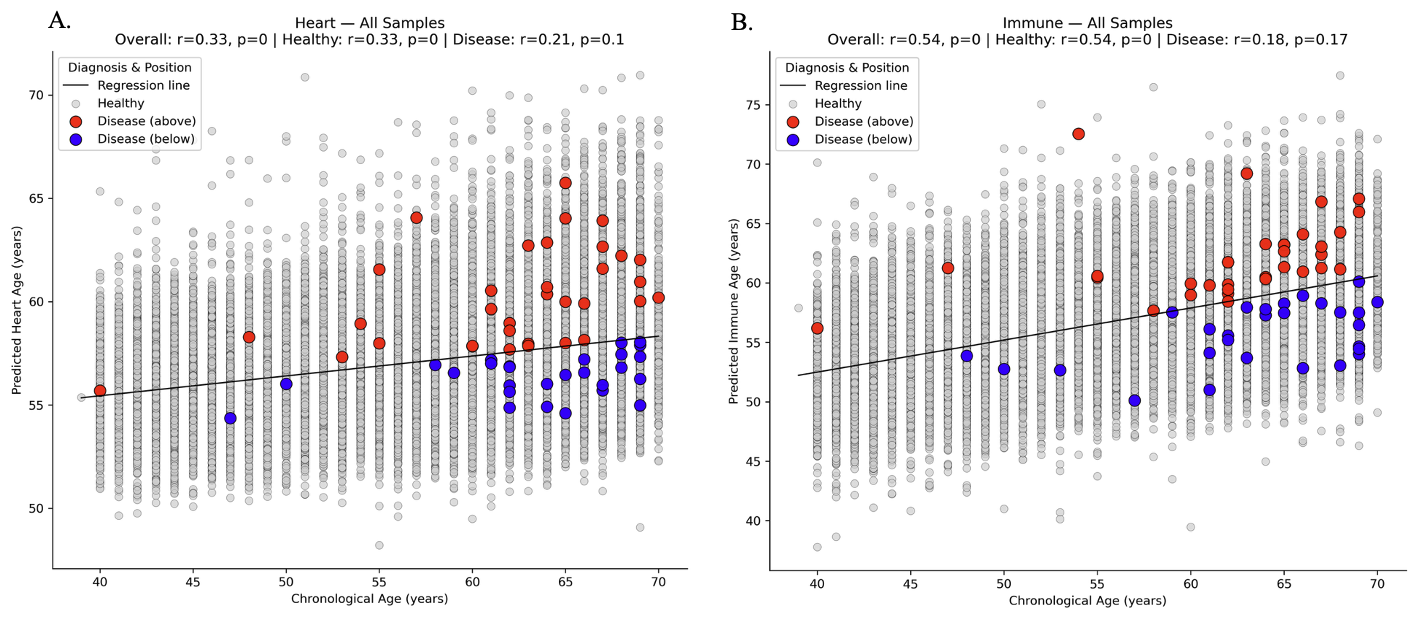

Figure S9. Re-analysis of proteomic age signatures in heart and immune ageing models, showing no clear acceleration relative to the UK Biobank reference trend. a. Heart ageing model. b. Immune ageing model.

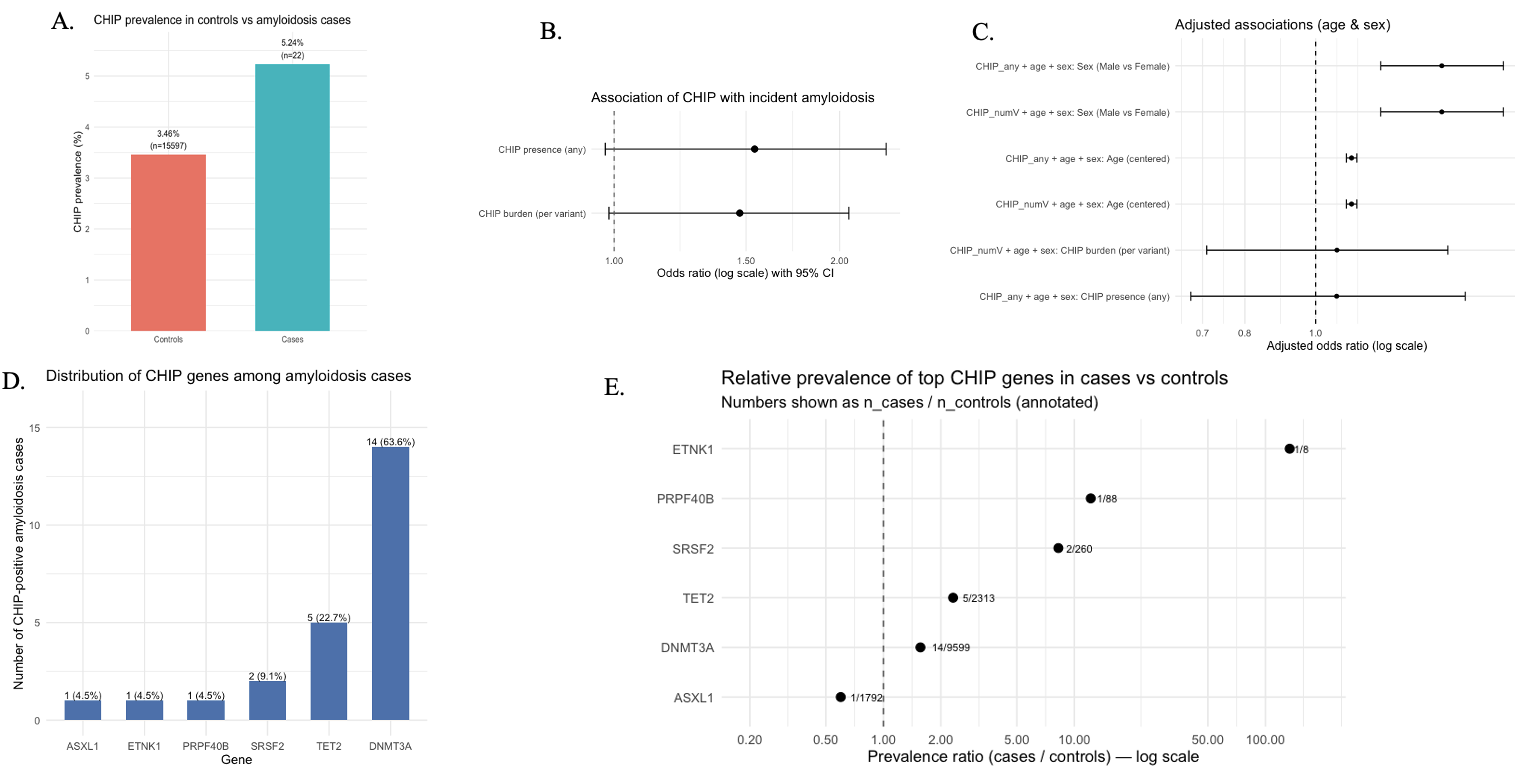

Figure S10. Clonal hematopoiesis of indeterminate potential (CHIP) analysis. a. CHIP prevalence in amyloidosis cases and controls. b. Unadjusted association between CHIP and incident amyloidosis (odds ratios with 95% CI). c. Age- and sex-adjusted associations showing attenuation of the CHIP–amyloidosis relationship. d. Distribution of CHIP-associated mutations among amyloidosis cases. e. Gene-level prevalence ratios (cases vs controls), highlighting variability driven by small sample sizes.

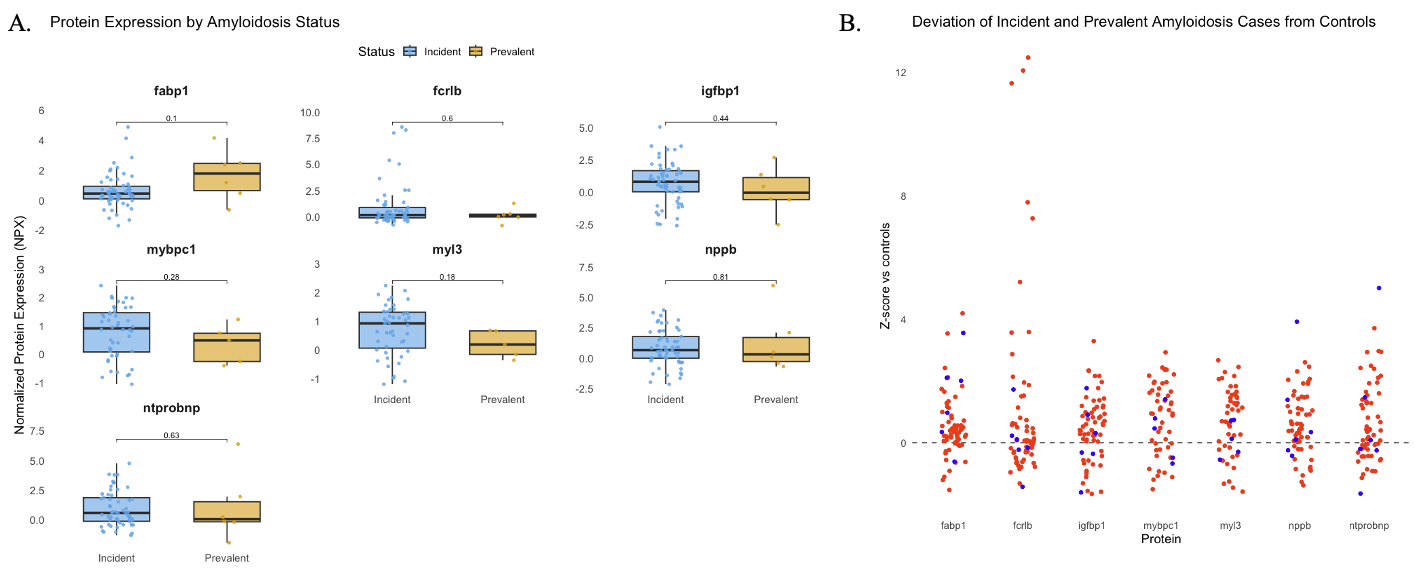

Figure S11. Comparison of key protein expression between prevalent and incident amyloidosis cases. a. Protein expression distributions by disease status. b. Standardised deviation from control levels across proteins.
